# Optimizing the Cell Painting assay for image-based profiling

**DOI:** 10.1101/2022.07.13.499171

**Authors:** Beth A. Cimini, Srinivas Niranj Chandrasekaran, Maria Kost-Alimova, Lisa Miller, Amy Goodale, Briana Fritchman, Patrick Byrne, Sakshi Garg, Nasim Jamali, David J. Logan, John B. Concannon, Charles-Hugues Lardeau, Elizabeth Mouchet, Shantanu Singh, Hamdah Shafqat Abbasi, Peter Aspesi, Justin D. Boyd, Tamara Gilbert, David Gnutt, Santosh Hariharan, Desiree Hernandez, Gisela Hormel, Karolina Juhani, Michelle Melanson, Lewis Mervin, Tiziana Monteverde, James E Pilling, Adam Skepner, Susanne E. Swalley, Anita Vrcic, Erin Weisbart, Guy Williams, Shan Yu, Bolek Zapiec, Anne E. Carpenter

## Abstract

In image-based profiling, software extracts thousands of morphological features of cells from multi-channel fluorescence microscopy images, yielding single-cell profiles that can be used for basic research and drug discovery. Powerful applications have been proven, including clustering chemical and genetic perturbations based on their similar morphological impact, identifying disease phenotypes by observing differences in profiles between healthy and diseased cells, and predicting assay outcomes using machine learning, among many others. Here we provide an updated protocol for the most popular assay for image-based profiling, Cell Painting. Introduced in 2013, it uses six stains imaged in five channels and labels eight diverse components of the cell: DNA, cytoplasmic RNA, nucleoli, actin, Golgi apparatus, plasma membrane, endoplasmic reticulum, and mitochondria. The original protocol was updated in 2016 based on several years’ experience running it at two sites, after optimizing it by visual stain quality. Here we describe the work of the Joint Undertaking for Morphological Profiling (JUMP) Cell Painting Consortium, aiming to improve upon the assay via quantitative optimization, based on the measured ability of the assay to detect morphological phenotypes and group similar perturbations together. We find that the assay gives very robust outputs despite a variety of changes to the protocol and that two vendors’ dyes work equivalently well. We present Cell Painting version 3, in which some steps are simplified and several stain concentrations can be reduced, saving costs. Cell culture and image acquisition take 1–2 weeks for a typically sized batch of 20 or fewer plates; feature extraction and data analysis take an additional 1–2 weeks.

**Key references using this protocol:** *Virtual screening for small-molecule pathway regulators by image-profile matching* (https://doi.org/10.1016/j.cels.2022.08.003) - recent work examining the ability to use collected Cell Painting profiles to screen for regulators of a number of diverse biological pathways.

*JUMP Cell Painting dataset: images and profiles from two billion cells perturbed by 140,000 chemical and genetic perturbations* (DOI) - the description of the main JUMP master public data set, using this protocol in the production of >200 TB of image data and >200 TB of measured profiles.

**Key data used in this protocol:** *Cell Painting, a high-content image-based assay for morphological profiling using multiplexed fluorescent dyes* (https://doi.org/10.1038/nprot.2016.105) - this paper provides the first step-by-step Cell Painting protocol ever released.

## Introduction

Since the advent of the digital camera, computationally-minded biologists and biologically-minded computationalists have realized that microscopy images represent an incredibly rich source of data; a camera that produces images that are 1,000 pixels on each side produces a million quantitative data points with every acquisition. The idea that scientific answers could be extracted from those pixels underlies the field of *image-based profiling* (also known as *morphological profiling*): its premise is that this pixel information, the details of which are often invisible to the human eye, are sufficient to cluster samples to draw meaningful conclusions.^1,2^

The Cell Painting assay (Figure 1) is a major driver of this field’s success - the creation of an assay that could be done simply and inexpensively on commonly available lab equipment provided researchers in both academia and industry the ability to try morphological profiling on their own data and an opportunity to standardize feature sets across laboratories. The two initial papers ^3,4^ describing the assay have been cited more than 500 times combined. Briefly, Cell Painting involves staining the cells the scientist wishes to characterize with 6 small-molecule dyes which together mark 8 organelles or cell compartments (DNA, cytoplasmic RNA, nucleoli, actin, Golgi apparatus, plasma membrane, endoplasmic reticulum, and mitochondria). Once the stained cells are imaged, three major compartments (whole cells, nuclei, and cytoplasm) are segmented using image analysis software and hundreds or thousands of image measurements (such as sizes, shapes, intensity distributions, and textures) are calculated from each cell. These measurements can then be used in a variety of ways, but one very common approach is to aggregate the single-cell data per well, perform normalization and feature reduction, then calculate the overall similarity in “feature space” between pairs of wells to search for unexpected or revelatory very high or very low similarity measurements. Cell Painting has been used to profile mutations in lung cancer ^5^, find treatments for COVID-19 ^6^, and help explore the toxicity of environmental chemicals ^7^ among dozens of other applications^1^. Scientists have also started creating their own variants, swapping out stains for lysosomes rather than mitochondria ^8^ or lipid droplets rather than the endoplasmic reticulum ^9^. We therefore believe that new stain combinations can be powerful for particular biological areas of interest, but we also recognize the value of a standardized assay performed in many laboratories and on many biological questions across the community in order to share data.

**Figure 1.**
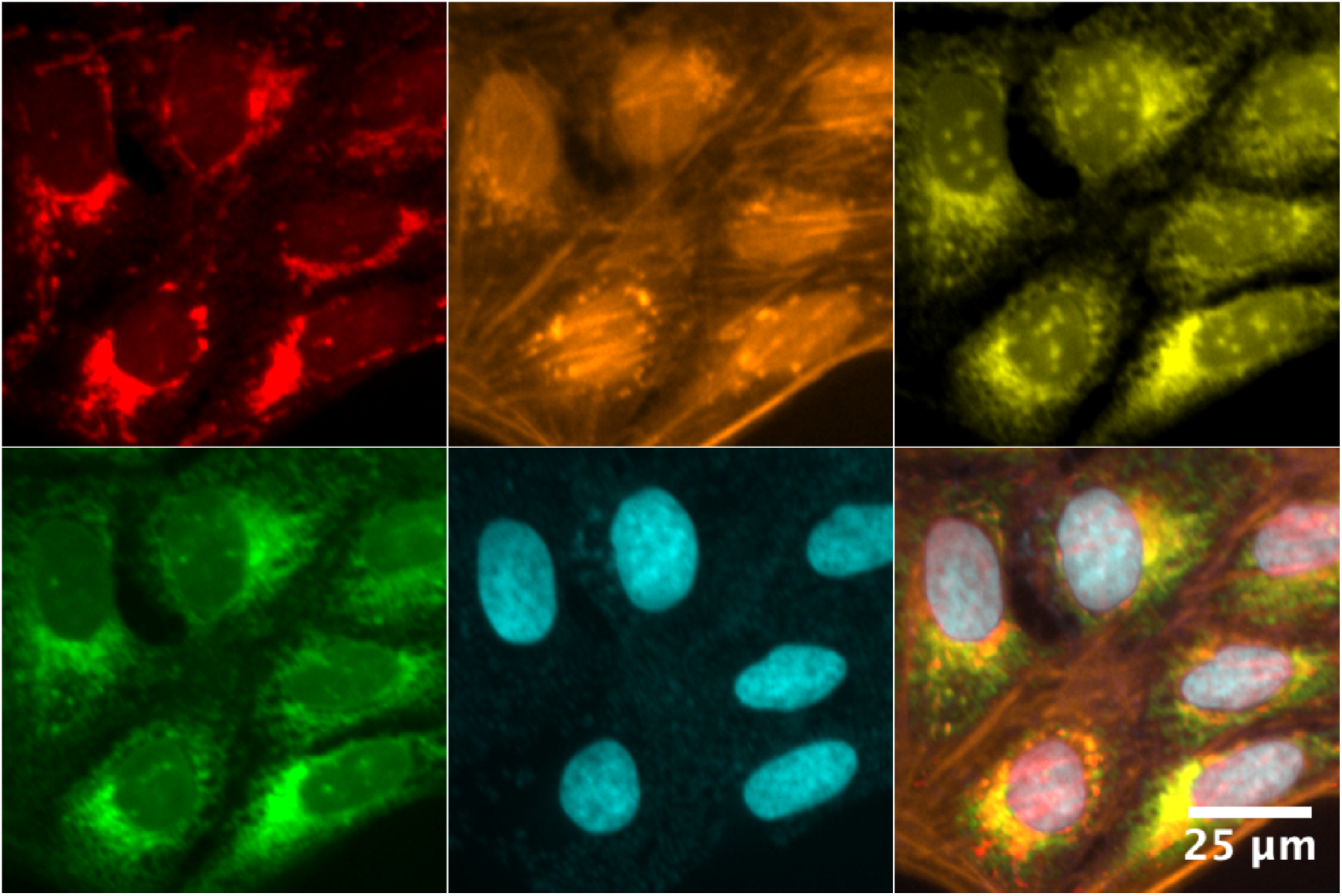
DMSO-treated U2OS cells treated with the Cell Painting assay; the six dyes in five channels stain eight cellular compartments. Top Row (L-R): Mitochondrial staining; Actin, Golgi, and Plasma Membrane staining; Nucleolar and cytoplasmic RNA staining. Bottom Row (L-R): Endoplasmic reticulum (ER) staining; DNA staining; a montage of all five channels.

We recently became interested in creating a very large public Cell Painting data set due to the success of multiple approaches where image data can be used to predict not just particular phenotypes about the parts of the cell that were stained, but entirely orthogonal data such as gene expression data or the outcome of a biochemical assay ^10,11^. Such a database could also allow queries for compounds that match genes ^12^ and vice versa. It stands to reason that a large, well constructed set of image-based phenotypic screen data could not only inspire new computational tools to mine such data, but also serve as a resource for researchers to compare their own data against, accelerating discovery for thousands of scientists globally. The Joint Undertaking for Morphological Profiling (JUMP) Cell Painting Consortium was formed in 2019 to create such a dataset, with a goal to optimize and select a single set of staining conditions prior to creating more than 150,000 perturbation profiles across a dozen sites around the world.

With these goals in mind, in this work we describe the optimizations done over more than a year to select optimal conditions for the JUMP Consortium’s production data set. More than two dozen different parameters were optimized, often testing several options per parameter and in the context of multiple combinations, yielding 299 staining-plate-imaging-runs or “logical plates”; a full matrix of every parameter would be greater than 50 million logical plates and as such was not attempted. Logistical constraints, and the fact that all parameters are deeply interconnected, prevented a fully iterative parameter testing strategy. A table describing all conditions tested is available as Supplementary Data 1.

Here, we present our updated recommendations, as well as our major finding: Cell Painting is remarkably robust. While most of our attempted optimizations admittedly did not dramatically change any parameter, the vast majority of conditions led to consistently similar results. In nearly all conditions, for our control compound plate, we observed a ***percent replicating*** (how often the profile similarity between two individual wells of Treatment X have pairwise similarity greater than expected; see below) of 70% +/-15%. Similarly, across nearly all conditions we observed a ***percent matching*** (if Treatment X and Treatment Y are thought to work via the same mechanism of action, how often are the profile similarities of a well of Treatment X and Treatment Y greater than expected; see below) of 20% +/-10%. This robustness did not yield an exciting optimization process, but speaks well of the stability of the assay and how likely it is to work across a variety of laboratories worldwide, even if a given laboratory may need to adjust some parameters for their local conditions. Here, we describe our optimization findings and decisions so that a new researcher can consider these questions when setting up their own Cell Painting experiments.

The major changes between our previous protocol and the current recommendations are as follows (presented in order of the protocol steps):

1. No media removal before the addition of MitoTracker, to simplify the protocol and minimize the loss of cells
2. Our recommendation for MitoTracker staining concentration remains at 500nM, but previous versions of the protocol used instructions that unintentionally led to a lower final concentration (375nM). The current protocol ensures a 500nM concentration after dilution.
3. Combining permeabilization and staining steps to make the process more automation friendly
4. Reduction of Phalloidin 4-fold, from 5µL/mL (33nM) to 1.25µL/mL (8.25nM), to save reagent costs
5. Reduction of Hoechst 5-fold, from 5µgmL to 1µg/mL, to save reagent costs
6. Increase of SYTO 14 2-fold, from 3µM to 6µM, to improve its signal
7. Reduction of Concanavalin A 20-fold, from 100µg/mL to 5µg/mL, to save reagent costs
8. Overall reduction of post-fixation staining volumes from 30µL/well to 20µL/well, to save reagent costs

### Optimization setup

A conventional assay is optimized by assessing each variant of the protocol for the optimal separation between a particular positive control and negative control. A profiling assay, by contrast, aims to measure hundreds to thousands of readouts, creating a challenge for optimization. Our prior efforts at optimization relied on assessing signal quality by eye ^3,4^. By contrast, here we aimed to make the first attempt at quantitative optimization. We evaluated each variant of the assay using 90 compounds, selected as detailed later, to cover a broad spectrum of biological activities.

We selected optimal parameter settings based on two metrics calculated on image-based profiles derived from Cell Painting cells treated with those compounds: ***percent replicating*** and ***percent matching***. Both describe how often a given pair of wells that should be similar actually are similar; specifically, how often is their pairwise correlation across features greater than the 95th percentile of a null distribution of the similarities of 10,000 pairs of random (non-matching) wells^13^. In percent replicating, the two wells that are compared are treated with the identical treatment and should perfectly correlate, if not for technical variations. In percent matching, the two wells are treated with different treatments that are believed (due to outside knowledge, i.e. ground truth) to produce similar biological impact. Percent replicating expects identical treatments to produce positive correlations, so it is scored in a one-tailed fashion (fraction of profile pairs above the 95th percentile of correlation values); by contrast, pairs of treatments might be expected to correlate or anti-correlate, so percent matching can be calculated as the fraction of treatments below the 5th percentile, the fraction above the 95th percentile, or both. These metrics have limitations, as elaborated previously ^14^, but were sufficient at the time for the purposes here: selecting the optimal assay conditions given the presence of replicates in different well positions on our compound control plate, and identical numbers of replicates of each condition. When comparing conditions in a given experiment, percent replicating has the advantage of being calculated for a higher number of conditions, because it is computed for each treatment individually. By contrast, percent matching has the advantage of better approximating an application of image-based profiling: matching different samples with each other, but because it can be calculated only for pairs of samples, there are fewer classes and thus less statistical power. It is also a harder task because most compounds annotated as having a target in common do not in fact produce similar morphological profiles, particularly due to polypharmacology ^13^ (see Supplemental Methods) and because ground truth annotations are incomplete and imperfect. Nevertheless, because samples that are supposed to match each other are in different positions within our compound control plate layout, this metric is usually less influenced by plate position effects, such as edge effects, which can unfairly improve percent replicating when replicates are in identical well positions within the plate layout.

The percent replicating and percent matching of an entirely untreated or negative control plate would be expected to be 5% (or 10%, if both tails of the distribution are assessed). Our first step was to select a set of standardized controls to determine whether treated wells were matching more than would be expected by chance. For initial optimizations, a compound plate known as the JUMP-MOA plate was created (Figure 2A, Extended Data Figure 1); in 384 wells, it contains 24 DMSO wells and 90 compounds at four replicates each. The 90 compounds are from 47 diverse mechanism-of-action (MOA) classes, with 43 classes having two compounds each and four classes having only a single compound. This allows for testing percent replicating for each of the 90 compounds and percent matching for each of the 43 multi-compound MOA classes within a single plate, allowing each plate to serve as its own “batch”; see Supplemental Methods for more information. This plate layout was used for most optimizations of staining reagents and conditions, imaging conditions, and feature measurements; percent matching for these plates is calculated in a one-tailed fashion (>95th percentile).

**Figure 2.**
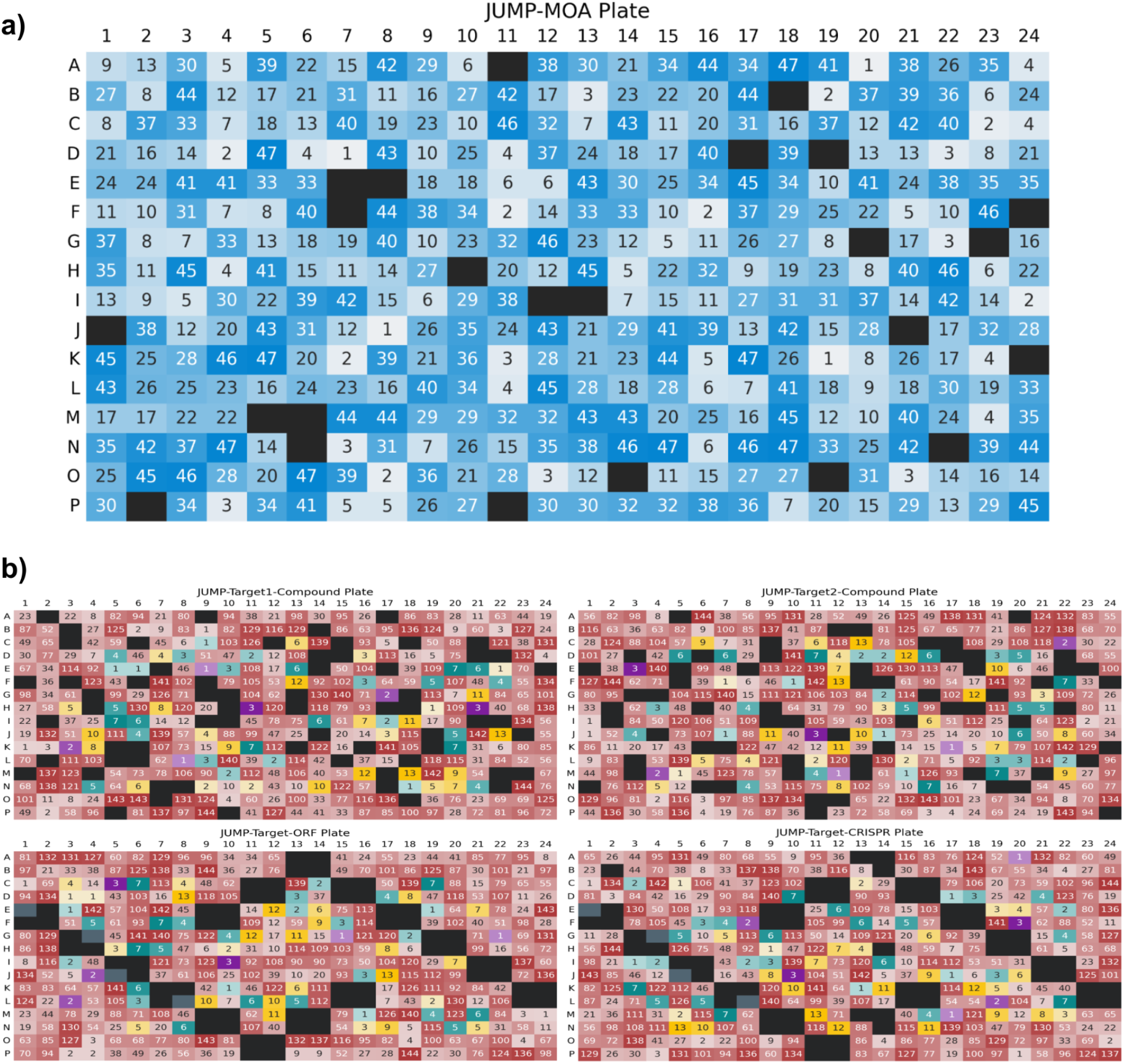
A) The JUMP-MOA compound plate map: unlabeled wells are DMSO only, all other wells are labeled to show distribution of MOA classes across the entire plate. A version of this figure grouped by compound rather than MOA is available as Extended Data Figure 1. B) The JUMP-Target plate maps. Black wells contain negative control treatments, while grey wells are untreated; other sets of control treatments were selected to either provide sets of diverse pairs of positive controls (purples), to provide a match between genes and compounds based on previous Cell Painting experiments (teals) or to match genes and compounds based on external reports of strong correlations between pairs (yellows). These controls are scattered among treatments hypothesized to affect other genes (reds).

To assess whether optimal conditions would differ across the different kinds of treatments planned for the JUMP consortium (compound treatments, open reading frame (ORF) overexpression, and CRISPR knockout), a second set of plates known as the JUMP-Target plates were produced (Figure 2B). These are described elsewhere^13^; briefly, they contain either compounds, ORFs, or CRISPR knockdowns related to a set of >175 genes thought to have strong and/or diverse phenotypic effects. Because this set of samples is so large, to reach the same four replicates per treatment found in JUMP-MOA, one must create four identical treated plates of the JUMP-Target-Compound source plate (there are two versions of this plate with the same set of 306 compounds but in different layouts: JUMP-Target-1-Compound and JUMP-Target-2-Compound, hereafter JUMP-Target1 and JUMP-Target2). The JUMP-Target-CRISPR source plate also requires four identical treated plates; it contains two guides for most genes, arrayed in separate wells on each plate. By contrast, the JUMP-Target-ORF plate has 130 ORFs duplicated on the plate (because typically only a single ORF is available per gene), and thus needs only two treatment plate copies of the source plate in order to get a sufficient number of wells with four replicates. As JUMP-Target treatments may be expected to increase or decrease the function of a target gene, percent matching is calculated in a two-tailed fashion (<5th percentile or >95th percentile).

### Optimization of cell line selection, treatment and culture conditions

The furthest-upstream task that the consortium needed to consider was which cell line to use in data production, given our desire to have all data in one cell line to maximize matching across treatments. In addition to typical imaging-assay concerns such as flatness, we wanted specifically to know for each candidate line: 1) how many diverse phenotypes could we detect using the Cell Painting Assay, 2) how well it overexpressed exogenously introduced genes, given our plans to analyze ORFs, and 3) how well particular Cas9-expressing clonal lines could knock down genes, given our plans to analyze CRISPR reagents. Especially when reproducibility across sites is critical, one additionally may wish to consider the availability of Cas9-expressing clones (some lines carry restrictive licenses making them unavailable to the public).

Once a line was chosen, we also wanted to know the effects of several variables in the Cell Painting assay protocol: 4) how long should a given treatment be applied to the cells, 5) how sensitive would the assay be to changes in plating density, 6) how if at all using Cas9-expressing clonal lines for CRISPR experiments and parental lines for compounds would affect our ability to match treatments affecting the same mechanisms of action, 7) how if at all using drug selection in our CRISPR and ORF conditions would affect their phenotypes and therefore our ability to match these conditions to compound treatment plates, and 8) whether or not cell conditions that some treatments would be exposed to (polybrene for improving lentiviral introduction of ORF and CRISPR reagents) should be applied to even the cells that did not *need* to experience them (such as the compound-treatment plates).

We focused on assessing the relative performance of A549 vs U2OS, because both are lines in which large public Cell Painting data sets already exist ^15^; dozens of other cell lines have performed well for Cell Painting experiments and researchers should choose a line that demonstrates phenotypes in which they are interested. We tested parental populations and several Cas9-expressing clonal lines for each cell type (one polyclonal of each line, with one additional monoclonal line for U2OS and three monoclonal lines for A549) using the JUMP-MOA plates to assess ability to detect multiple phenotypes (Figure 3A); Cas9 editing efficiency of the Cas9 expressing lines was measured in parallel (Supplementary Table 1, Supplementary Methods). In U2OS, the polyclonal line displayed much greater CRISPR efficiency as well as better percent matching. In A549, any if not all of the lines could have been suitable based on efficiency and percent matching; we therefore chose based on the line with the fewest restrictions on sharing between partners, which for the particular four clones in question was the polyclonal line. Additional experiments (quantified by cell count only) helped decide final cell densities, viral amounts, and other conditions (Supplementary Methods)

**Figure 3.**
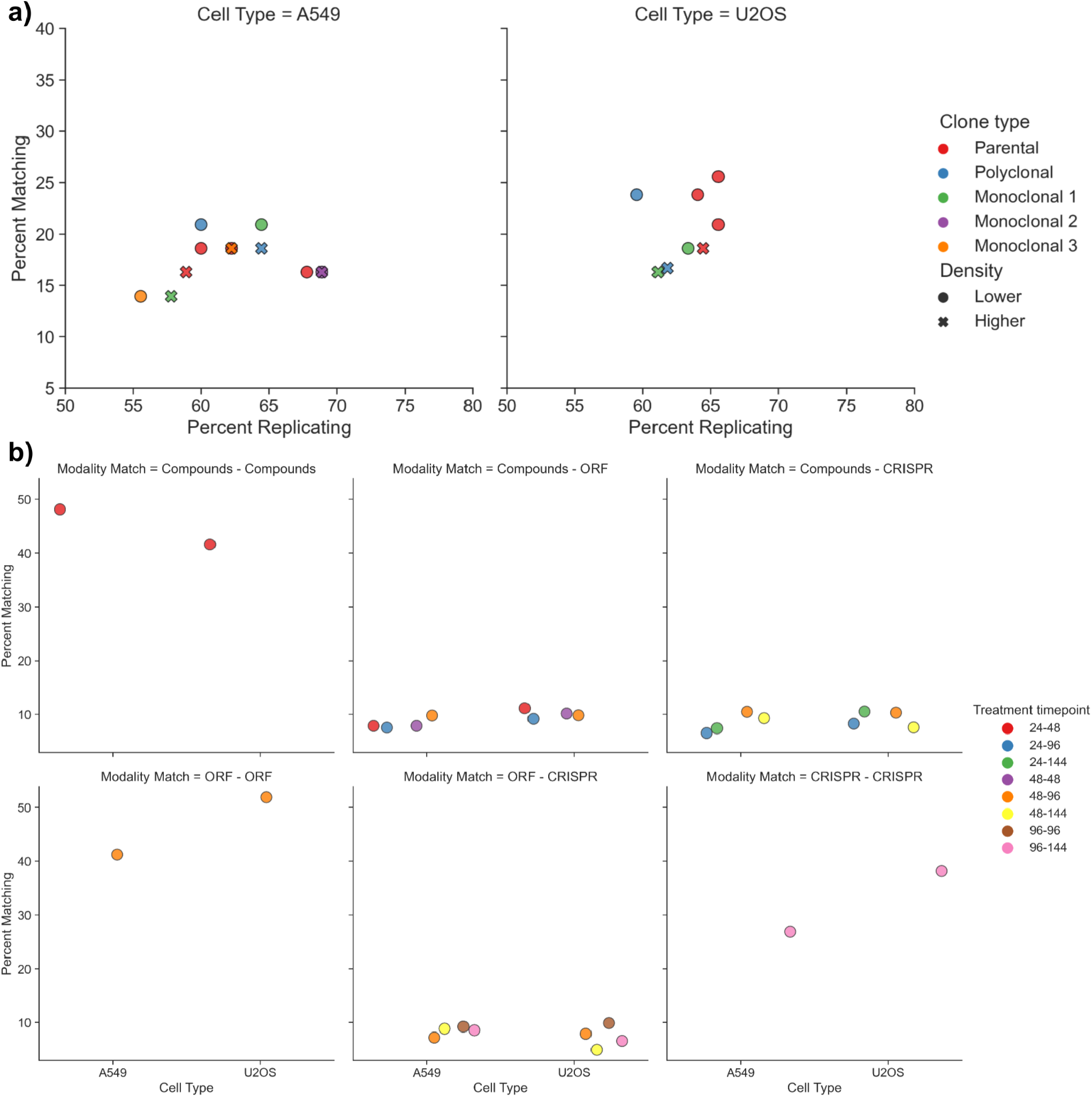
A) Assessment of compound percent replicating and percent matching using JUMP-MOA plates in A549 and U2OS parental, Cas9-polyclonal, and Cas9-monoclonal lines. In U2OS, the polyclonal line outperforms the single monoclonal line tested; in A549, performance is more varied. B) U2OS has slightly worse compound-to-compound-across-time-point percent matching than A549 but otherwise generally performs slightly better at percent matching when assessed via the JUMP-Target plates.

We next carried the parental and polyclonal lines forward to a single large experiment that we called CPJUMP1^13^, using the JUMP-Target plates to address questions 1-7 above; question 8 was assessed in a second, later experiment. Neither line performed poorly in any of our experiments, and on the whole none of the cross-modality matching performed much beyond what would be randomly expected (10%). On the whole U2OS displayed a slightly higher cross-modality matching and higher genetic perturbation-across-timepoint matching (Figure 3B), as well as higher within-plate compound matching (Figure 3A), and had a greater amount of publicly available data; thus by a slim margin U2OS was selected to be used for producing the JUMP-CP Consortium’s full dataset (questions 1-3). Based on these results, we also decided that compound treatments would be run at 48 hours of treatment, ORF overexpression experiments at 48 hours of treatment, and CRISPR knockdowns at 144 hours of treatment (question 4, Figure 3B).

We further analyzed the rest of CPJUMP1 to address questions 5-6; while these analyses were done in A549 rather than U2OS, we believe the results are likely to hold in both types due to their generally similar behavior. We saw little effect of profile sensitivity to plating density changes of +/-20% (Extended Data Figure 2A). We also saw little difference on modality matching when running compound experiments in the parental line vs the polyclonal Cas9 line (Extended Data Figure 2B), and as such chose to have all partners running compound experiments do so in the commercially available parental U2OS lines.

To answer question 7, we needed to assess for our ORF and CRISPR conditions how selection of the lentiviral reagents by resistance markers in the viral backbones would affect our ability to match between those conditions and our larger compound panel. Selection could possibly alter the results in either direction, improving them by removing cells not expressing the treatment and/or harming them by introducing a second “drug selection” signature that might perturb biological signals. Drug selection may have had a small deleterious effect on percent replicating (Extended Data Figure 3B) though we cannot rule out that this is due to fewer replicates for the selected plates than the unselected ones, and our assessment of cross-modality percent matching suggests a potential small deleterious effect from drug selection, especially in ORFs (Extended Data Figure 3A). This is understandable since in our case ORFs have a vector with a slower selectable marker (blasticidin) than CRISPR (puromycin) and are also treated for a shorter period of time (96 hrs vs 144 hrs). We therefore chose not to perform drug selection in our final protocol recommendations.

Finally, for question 8 we wanted to know whether we should include polybrene, which is used in aiding lentiviral transduction in our ORF and CRISPR plates, in our compound treatment production plates; as with the question of drug selection, one could *a priori* imagine it either helping or harming cross-modality matching. In this case we saw a strong deleterious effect on profiling results from a 24 hour treatment with 4 µM polybrene, with cross-plate-replication between polybrene-treated vs untreated plates dramatically lower than the intra-treatment cross-plate-replication of either treated or untreated plates (Extended Data Figure 3C). Polybrene addition also did not improve the ability to match Target2 plates to a previous batch of ORF overexpression plates (Extended Data Figure 3D), causing us to recommend against including it.

### Optimization of plates, staining reagents and conditions

Once a researcher has picked their cell line and treatment conditions, the next thing they must do is get their cells onto imaging plates and stain them. We saw no consistent difference in percent replicating or percent matching between plates from two manufacturers, one of which contained “barrier wells”, aka reservoirs at the edges of the plates that hold liquid in them to try to create even humidification of the whole plate surface (Extended Data Figure 4A). They may prove beneficial in other experimental contexts or research environments, especially those new to running sensitive high content assays like Cell Painting, but we decided against using them due to increased cost.

In order to try to minimize disruption to the cells and time and buffer spent washing plates, we have introduced two washing-related changes in this protocol vs our 2016 recommendations. First is to not remove any media from the wells before the addition of MitoTracker; this creates higher costs for this particular reagent, because staining is done in a larger volume, but it omits a step (removing the medium) and decreases the likelihood of precious cells becoming detached before fixation.

Second, the original protocol involved permeabilizing the cells, washing them twice, and then adding the other (non-MitoTracker) dyes; now we recommend performing the permeabilization and staining simultaneously. This leads to a shorter and easier protocol, and with no apparent negative consequences to profile quality (Extended Data Figure 4B). While the total volume for the MitoTracker staining step has gone up from 30µL to 60µL, all other dyes are now added in a smaller volume (20µL rather than 30µL).

The next thing a researcher setting up a Cell Painting assay must decide is exactly which stains to use and in which concentrations. Although variations on Cell Painting can be powerful for investigating particular areas of biology ^8,9^, for optimization of the canonical assay for this public dataset, the consortium only considered the original six dyes used in earlier versions of the assay (with the exception of DAPI for Hoechst, see Extended Data Figure 5B). We did briefly qualitatively assess if there were any benefits of moving WGA to an ultraviolet dye or MitoTracker to an orange dye, but quickly dismissed them: no immediate large improvement was shown (data not shown) and it would require reoptimization of the entire stain panel. A large number of stain concentration adjustments were tested; they are broken out comprehensively in Extended Data Figure 5. We found that reducing Hoechst from its original 5µg/mL to 1µg/mL and diluting Phalloidin from 33nM to 8.25nM had no ill effect and possibly even a positive one (Extended Data Figure 5B), so we adopted these changes to reduce reagent costs and waste.

All of the changes in the preceding paragraphs form what we call the Cell Painting version 2.5 protocol; this protocol was used for the CPJUMP1 experiment and represents an improvement in percent replicating and percent matching from the version 2 protocol from 2016 ^3^ (Figure 4A). However, some consortium members reported higher than optimal bleedthrough in version 2.5, specifically that on their microscopes there was so much crosstalk between the ER (Concanavalin A) and RNA (SYTO 14) stains that it affected detection of the RNA signal. We therefore did one additional round of optimization to yield the version 3 (v3) protocol, incorporating all of the improvements from v2.5 plus reducing 20x the amount of Concanavalin A (the most expensive dye in the panel) and doubling the amount of SYTO14 (Figure 4A). These changes brought the two channels better into balance and became our final recommendation, version 3 (v3). There are two major sources for the complete set of Cell Painting dyes-ThermoFisher and PerkinElmer. We tested both dye sets in a number of batches and across a variety of conditions and found their performance to be equivalent (Figure 4B).

**Figure 4.**
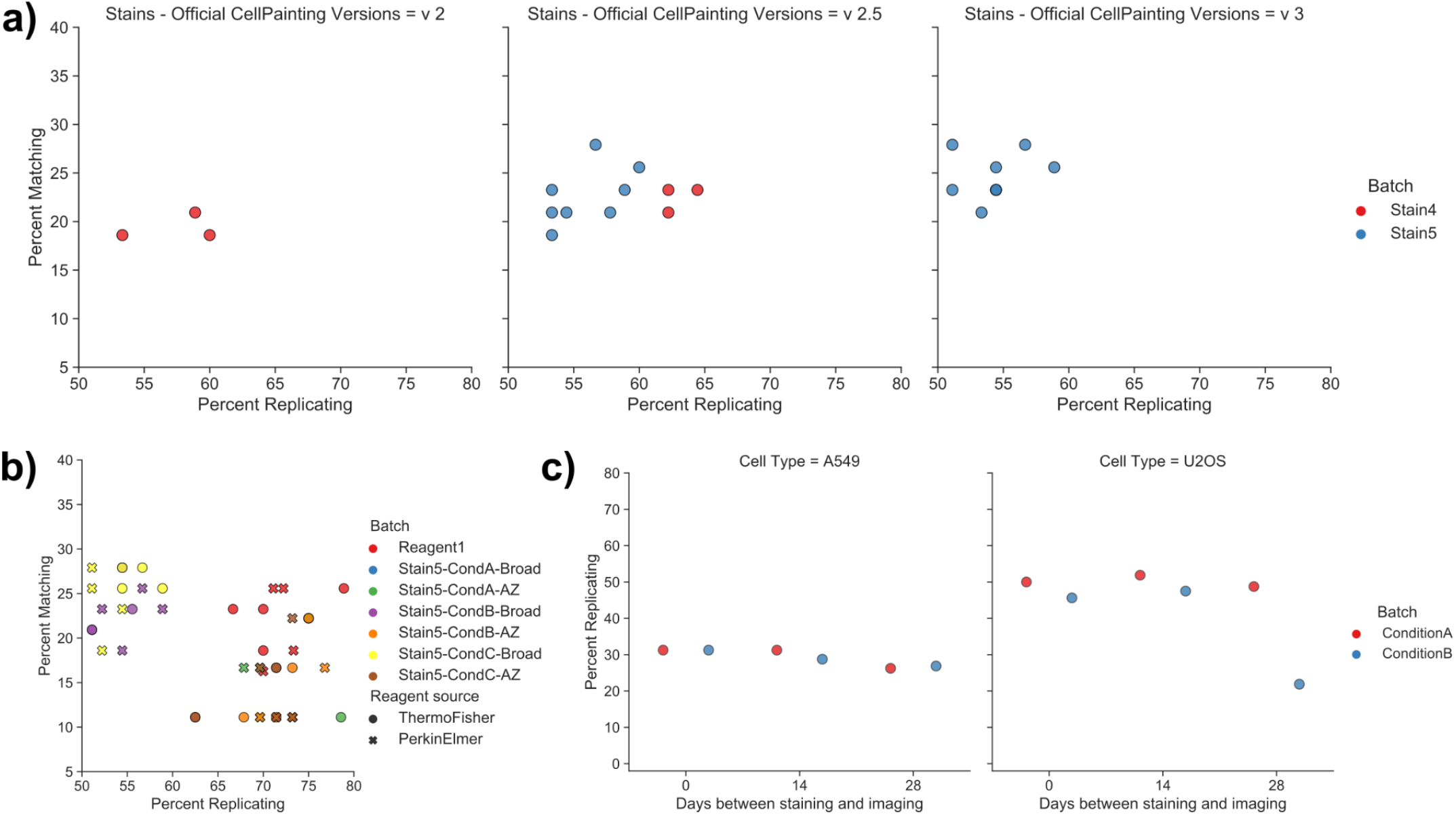
A) Comparison of percent replicating and percent matching for 3 versions of Cell Painting - version 2 (Bray et al 2016 conditions) vs version 2.5 (Bray et al plus introduction changes 1-6) vs version 3 (this manuscript’s final recommendations). The move from v2 to v2.5 seemed to improve both percent replicating and percent matching in the Stain 4 pilot experiment (red dots); the move from v2.5 to v3 decreases reagent cost while maintaining comparable if not slightly improved percent matching in the Stain5 pilot experiment (blue dots). Note that Stain 5 experiments (blue dots) were performed using only half of the compound dose as Stain 4 experiments (red dots) B) Comparison of reagents from two different vendors across multiple stain conditions and microscopes. Performance is extremely similar between vendors in all conditions tested. C) Assessment of persistence of Cell Painting plate quality over storage time. Percent replicating seems to be decreasing by Day 28 but is quite similar to initial values at day 14.

The Cell Painting assay uses fixed cells, reducing the need for precise timing of the image acquisition step. Because microscopes occasionally break or are booked, we finally wanted to test the timescale of deterioration for a fixed and stained Cell Painting plate, in terms of overall profile quality; we see no measurable degradation of profile percent replicating between plates imaged on the day they were stained and plates imaged 14 days later. There appears to be a consistent decline between week 2 and week 4, however, so if possible we recommend imaging is completed within 28 days after staining (Figure 4C).

### Optimization of imaging conditions

Once the staining conditions are finalized, the next step is to optimize the image acquisition conditions. As in other areas of optimization, we had a number of questions - 1) Which kind of scope should we use? 2) Should we take one Z plane or several? 3) Should the camera be set to 1×1 or 2×2 pixel binning? 4) How high should our exposures be? 5) How many fields of view do we need to take? and 6) If a researcher needs to image a plate more than once (during optimization or due to a technical failure), how much deterioration of signal can they expect?

In our testing, the answer to most of these questions was simply “probably anything will produce comparable results”. We saw no consistent difference between images taken in widefield vs confocal (Figure 5A), even when those modes were on two entirely different microscopes (Extended Data Figure 6A). The only plate in which we saw decreased performance for confocal microscopy was in a condition where the microscope could not create a filter set match that would separate the ER and RNA data and thus captured them only as a single channel. To our surprise, there was essentially no profile quality loss in our hands by switching from capturing images with 1×1 binning to 2×2 binning (Figure 5C), leading us to choose 2×2 binning, because it reduces data storage price 4X and compute costs significantly. We also did not see any change between lower and higher exposures for each of several staining conditions (Figure 5D), suggesting that as long as the exposure times are set reasonably enough to maximize dynamic range while minimizing saturation, the exact values are less important and one can potentially save imaging time by using slightly shorter exposures.

**Figure 5.**
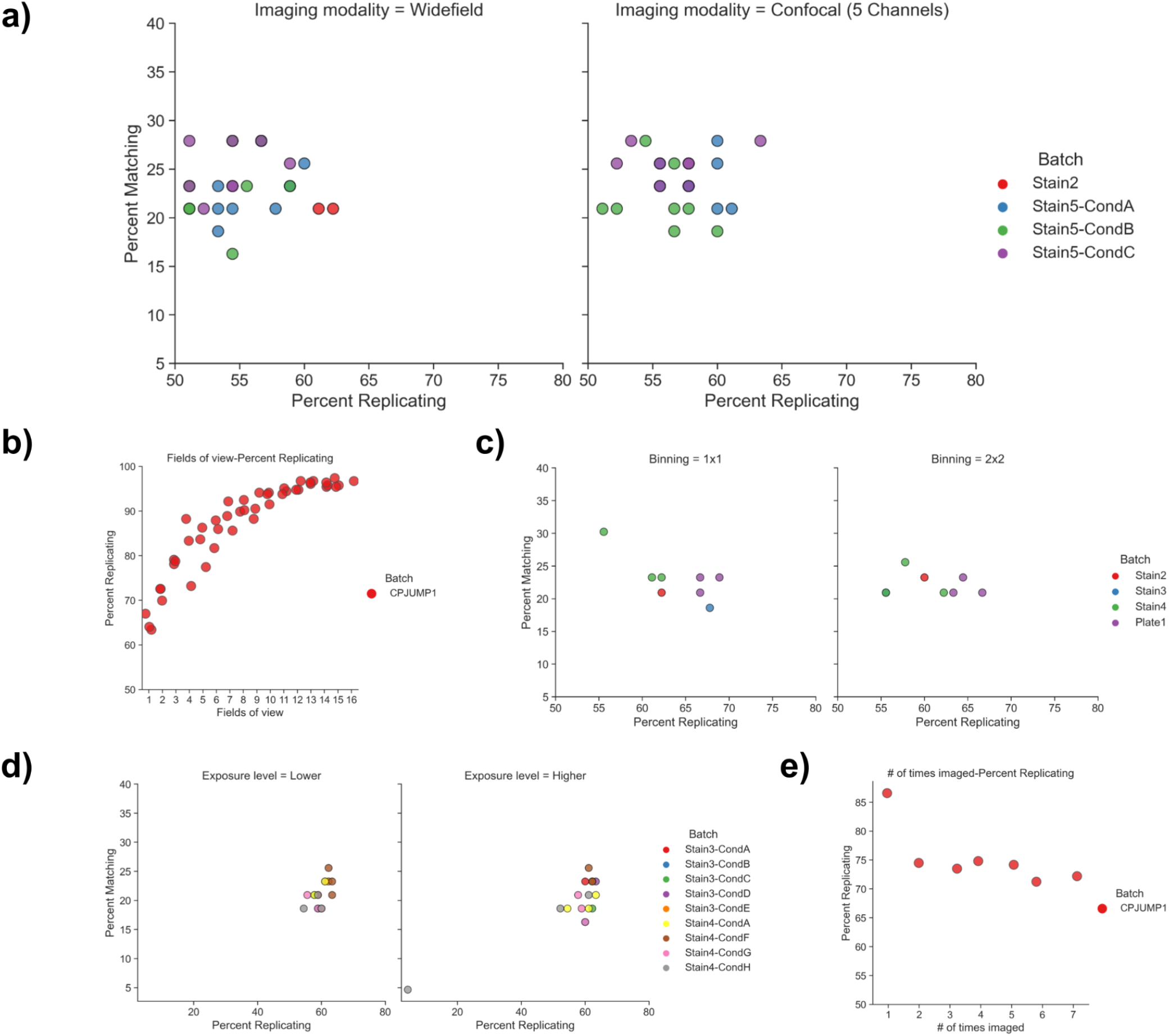
A) Cell Painting works similarly well with widefield and confocal microscopy. B) Relationship between fields of view captured and percent replicating - increasing the number of fields of view increases percent replicating up until approximately 10 fields of view. At this magnification and plating density, each field of view contains approximately 145 cells. C) Cell Painting performs well when images are captured with either 1×1 or 2×2 binning. D) Cell Painting performs similarly when plates are imaged at a lower or 2-4X higher (but still below saturation) laser power and/or exposure time. E) Assessment of effect of re-imaging on percent replicating: a drop between first and second imaging is observed and a potential small continued decrease thereafter.

A few parameters do seem to produce measurable results; we saw small increases in percent replicating (but not percent matching) in two experiments where we tested imaging the same plate in one vs several fluorescent Z planes. The increase was small enough (Extended Data Figure 6B) that we chose not to complicate our workflow but may be worth considering in other circumstances, such as when using confocal imaging or when using cells that have varying Z heights such as neurons. We also found that the number of fields of view acquired was important, with each additional field of view leading to an increase in percent replicating out to at least 10 fields of view (Figure 5B) at our magnification and plating density (approximately 145 cells per field of view). We suspect this has more to do with the total number of cells imaged than the number of fields of view *per se*. We saw a roughly 10% drop in percent replicating signal when plates had been imaged twice versus a single time, and minor subsequent losses thereafter up to six (Figure 5E). We therefore recommend that plates are imaged only once, though 10% loss may be acceptable in many contexts, relative to the cost of repeating sample preparation for a given plate or batch; as the initial percent replicating value was extremely high, a 10% loss is likely a worst-case scenario.

### Optimization of image analysis conditions

In our experience, creating accurate segmentations is crucial but difficult to optimize in a rule-driven way without substantial effort to create ‘ground truth’ for evaluating changes in the pipeline’s parameters ^16^. We have described elsewhere ^17^ a number of resources for learning to do so effectively, and the workflow described here includes steps for iterating on segmentation to ensure it is optimal before feature extraction. As deep learning segmentation tools such as StarDist ^18^ and Cellpose ^19^ become more popular, they may help solve many difficult segmentation issues. These tools may be used independently to create segmentations and then object labels matrices can be brought into CellProfiler, used via their ImageJ implementations using the RunImageJMacro ^20^ or with their CellProfiler plugins (https://github.com/CellProfiler/CellProfiler-plugins) if CellProfiler is installed from source. ^16^

Once objects are segmented, in general we recommend measuring as many image-based features as is practical in your image analysis software, where “practical’’ has a few considerations, such as a) the amount of time needed to generate these measurements and b) limitation on your output size based on file constraints - SQLite unless manually compiled only allows 2000 columns in a given table, for example.

An open question about Cell Painting is how much each stain contributes to the information content of the assay. Likewise, the contributions of different measurement categories are unknown. Therefore we analyzed the relative contributions towards percent replicating of different measurement types (Figure 6A), cell compartments (Figure 6B), and channels (Extended Data Figure 7) in the context of the 90 compounds present in the JUMP-MOA plate. As with the staining and imaging, for these phenotypes the data we collect is extremely robust to changes in the measured features and channels - most subsamples of channels and/or features and/or compartments (Figure 6A-B, Extended Data Figure 8, and Extended Data Figure 9) will still lead to a high-quality final analysis. We note however that these breakdowns describe only the phenotypes present in the JUMP-MOA plates; any specific phenotype(s) of interest may crucially depend on compartments, stains, or features that are less critical for these 47 mechanisms of action, and as such we always recommend capturing as many features as practicably possible. We saw only small differences when the scales of several CellProfiler features were adjusted, indicating that there is at least a reasonable range of tolerances for these parameters as well (Figure 6C).

**Figure 6.**
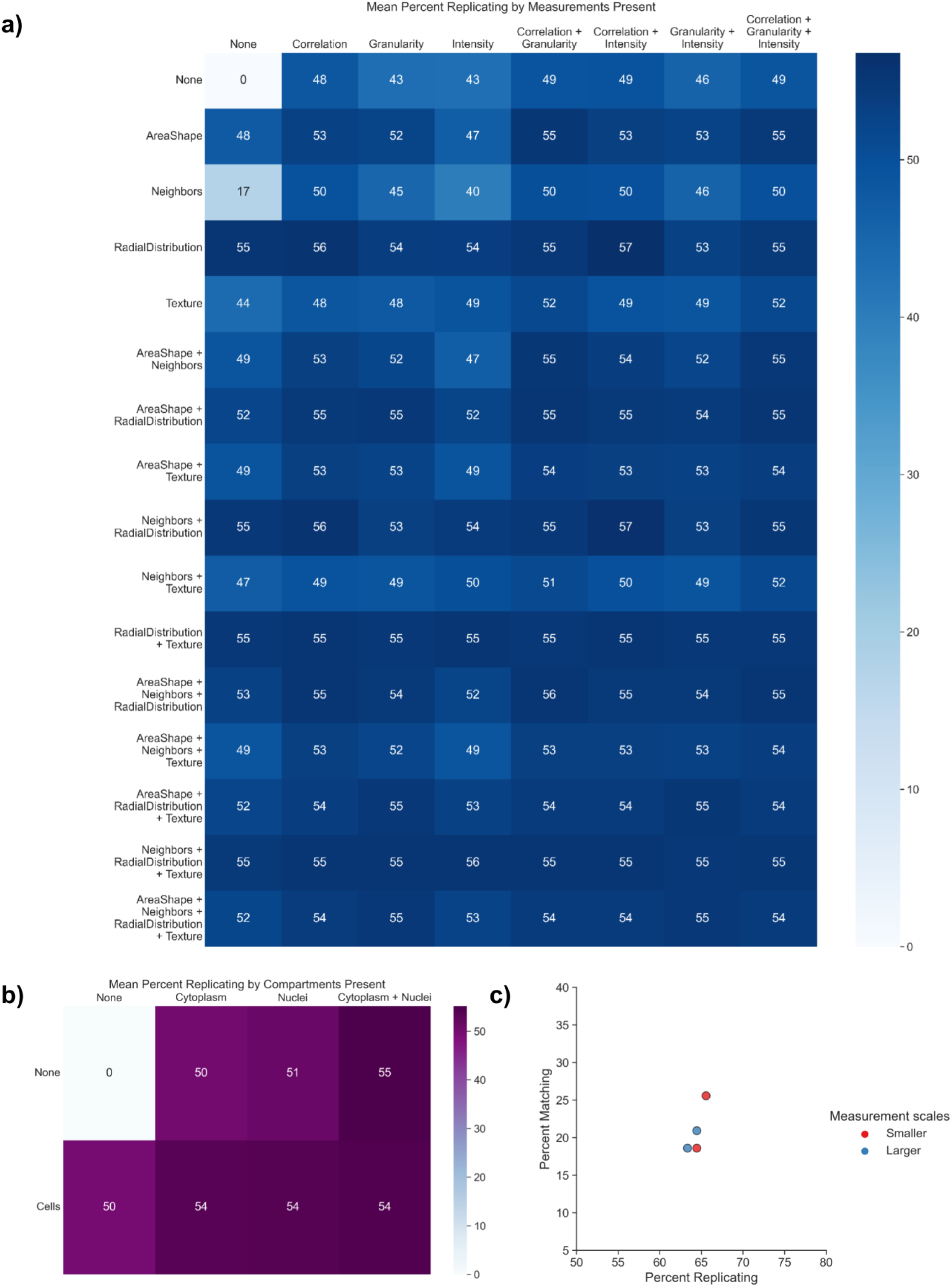
A) Mean percent replicating of eight JUMP-MOA plates stained with the final staining conditions after dropping out all possible combinations of features from the seven major feature categories before performing feature selection and calculation of percent replicating. To create a sufficiently compact data representation, the 7 categories present were split 3 onto the X (Correlation, Granularity, and Intensity) and 4 onto the Y (AreaShape, Neighbors, RadialDistribution, and Texture) axes; this allows visualization of the 127 possible unique combinations. A channel-by-channel breakdown of the importance of the feature categories is provided as Extended Data Figure 9. B) A parallel analysis of the same experiment as in part A, but with the 3 compartments present, 2 on the X axis (Cytoplasm and Nuclei) and Cells on the Y axis. C) Assessment of the effect of varying the measurement scales in CellProfiler (ie measuring Texture at 3 and 5 pixel spacings vs 5 and 10 pixel spacings) on percent replicating for two plates of the CPJUMP1 experiment. Using larger measurement scales in the MeasureGranularity, MeasureTexture, and MeasureObjectNeighbors modules seemed to produce a very small decrease in percent replicating but no major effect.

## Limitations

As with any assay, phenotypic or otherwise, Cell Painting will not be able to detect every phenotype - previous work found phenotypes for 50% of overexpressed genes^21^, and the percent replicating values found in this work suggest even in our handpicked treatments we only detect about 70% of phenotypes. We hope that this work will help guide researchers who want to adapt Cell Painting to a new system or create variants of Cell Painting that will definitely detect their phenotypes of interest to understand the necessary experiments needed to find the right conditions for their system. We also hope that the control plates designed for this work can make optimizations for future researchers.

In creating such a large public data set and making it easier to reproduce our exact technique, our goal is to help researchers better match the public set to their own samples of interest. However, major work still remains in being able to match samples across different experiments^12^; a major goal of the JUMP Consortium is to develop such methods. We hope future technical advancements will make it possible to match samples across widely variant assays, but until then, matching our protocol to the degree possible represents the best chance of being able to match to the public set. Future technical advancements may make it possible to predict the results of this specific assay from brightfield images^22^, but it is not yet clear how transferable assays trained on data made at one location will be on data from another location.

## Materials

### Reagents

- Cell line of interest—e.g., we have previously used U2OS cells (American Type Culture Collection (ATCC), cat. no. HTB-96, https://scicrunch.org/resolver/RRID:CVCL_0042) or A549 cells (ATCC, cat. no. CCL-185, https://scicrunch.org/resolver/RRID:CVCL_0023) for the majority of our large screens. We are also aware of successful Cell Painting experiments using MCF-7, 3T3, ES2, HCC44, HTB-9, HeLa, HepG2, HEKTE, SH-SY5Y, HUVEC, HMVEC, Ocy454, primary human fibroblasts, primary human hepatocytes, primary human adipose-derived mesenchymal stem cells, and primary human hepatocyte/3T3-J2 fibroblast cocultures.

**Caution** Cell lines should be regularly checked to ensure that they are authentic and that they are not infected with Mycoplasma.

- DMEM media (Corning, cat. no. 10-013-CV)
- McCoy’s 5A media (Life Technologies, cat. no. 16600108)
- FBS (Sigma-Aldrich Inc., cat. no. F2442-500ML)
- 0.25% Trypsin-EDTA (Corning, 25-053-CI)
- PBS (Sigma-Aldrich Inc., cat. no. D8537-6×500ML)
- Pen-strep (Life Technologies, cat. no. 15140163)
- Ethanol (Decon Labs, cat. no. V1401)
- Polybrene (Sigma-Aldrich Inc, cat. no. 28728-55-4)
- Blasticidin S HCl 10 mg/mL (Life Technologies. cat. no. A1113903)
- Puromycin (Sigma-Aldrich Inc., cat. no. P9620-10ML)
- (Optional) Small-molecule libraries, typically 10 mM stock in DMSO (e.g., Chembridge library or Maybridge library; the compounds from the JUMP-Target-1 and JUMP-MOA plates are available as Pre-Plated Cell Painting libraries from Specs https://www.specs.net/index.php?page=2019041215290210#preplatedsets)
- (Optional) 384w ORF lentivirus (recommended vector: pLX_304)
- (Optional) 384w CRISPR lentivirus (recommended vector: pXPR_003)
- (Optional) Compounds for spike in controls on viral plates:
  - AMG-900, 10 mM in DMSO (Selleckchem, cat. no. S2719)
  - LY2109761, 10 mM in DMSO (Selleckchem, cat. no. S2704)
  - Quinidine, 10 mM in DMSO (MedChemExpress, cat. no. HY-B1751)
  - TC-S 7004 (Tocris, cat. no. 5088/10)
- (Optional) CellTiter-Glo (Promega, cat. no. G7573)
- Dyes - either as a kit or individually; ThermoFisher reagents as in our prior version of the protocol ^3^ performed comparably (Figure 4B)
  - PhenoVue™ Cell Painting Kit (PerkinElmer, cat. no. PING22)
  - Individual dyes:
    - MitoTracker Deep Red (Invitrogen, cat. no. M22426) OR PhenoVue 641 Mitochondrial Stain (PerkinElmer, cat. no. CP3D1)
    - Wheat-germ agglutinin/Alexa Fluor 555 conjugate (Invitrogen, cat. no. W32464) OR PhenoVue Fluor 555 - WGA (PerkinElmer, cat. no. CP15551)
    - Phalloidin/Alexa Fluor 568 conjugate (Invitrogen, cat. no. A12380) OR PhenoVue Fluor 568 - Phalloidin (PerkinElmer, cat. no. CP25681)
    - Concanavalin A/Alexa Fluor 488 conjugate (Invitrogen, cat. no. C11252) OR PhenoVue Fluor 488 Concanavalin A (PerkinElmer, cat. no. CP94881)
    - Hoechst 33342 (Invitrogen, cat. no. H3570) OR PhenoVue Hoechst 33342 (PerkinElmer, cat. no. CP71)
    - SYTO 14 green fluorescent nucleic acid stain (Invitrogen, cat. no. S7576) OR PhenoVue 512 Nucleic Acid Stain (PerkinElmer, cat. no. CP61)
    - PhenoVue Dye Diluent A 5x (PVDDA1)
- 32% (wt/vol) Paraformaldehyde (PFA), methanol free (Electron Microscopy Sciences, cat. no. 15740-S)
- HBSS (10×; Invitrogen, cat. no. 14065-056)
- Triton X-100 (Sigma-Aldrich, cat. no. T9284)
- DMSO (Millipore Sigma, cat. no. D5879-100ML)

### Additional Reagents Needed for Thermo Protocol

- Sodium bicarbonate (Sigma-Aldrich, cat. no. S571)
- Methanol (Honeywell, cat. no. 34966)
- BSA (Sigma-Aldrich, cat. no. 05470)

**CRITICAL** We have performed the Cell Painting assay using the specific catalog numbers listed here. If you are planning on changing to a different product or vendor for a given reagent, reoptimization of that reagent for the protocol may be necessary.

### Equipment

Microplates: PhenoPlate (previously Cell Carrier-384 Ultra) Microplates, tissue culture treated, black, 384-well with lid, Part Number: 6057300. Other microplates that are compatible with the microscope will suffice, as long as they are validated for use in high-content imaging (typically plates with minimal offset from the plate “skirt” to the well bottom surface (e.g. “Low Base or “Ultra-Low Base”).

**CRITICAL** step: order all 384w plates from same lot

- 384-well, tissue culture treated, white plate with clear flat bottom, with lid, with barcode label (Corning, cat. no. 8793BC)
- T-175 culture vessel (Greiner bio-one, cat. no. 661160)
- 50 mL Falcon centrifuge tubes (Corning, cat. no. 352070)
- 250 mL Polypropylene Centrifuge Tubes with Plug Seal Cap (Corning, cat. no. 430776)
- Serological pipettes, various sizes (Greiner bio-one)
- Aluminum single-tab foil, standard size (USA Scientific, cat. no. 2938-4100)
- evaporation resistant microplate lids or gas permeable adhesive plate seals
- Tissue Culture Microscope (Olympus, model CKX41SF)
- Automated cell counter: Beckman Z1 Particle Counter (Beckman Coulter, Model Z1)
- Cytomat 5C tissue culture incubator at 37 °C, 5% CO2 (Thermo Fisher Scientific, cat. no. 50128822) or Liconic Instruments: Liconic incubator, Model: STX-220 HR (with stacker and rotating plate carousel) or Thermo Heracell VIOS 160i CO2 Incubator at 37 °C, 5% CO2 (ThermoFisher Scientific, Model Vios 160i)
- Automated liquid handler: Multidrop Combi Reagent Dispenser (Thermo Fisher Scientific, cat. no. 5840300) or Freedom EVO with 384-channel arm (Tecan, cat no. MCA384) or Janus (PerkinElmer, Model JANUSMPD). Pressurized valve dispensing systems (GNF WDII & OTD) are more consistent than peristaltic (Combi/EL406).
- 30 µLFilter Tips, Sterile for Janus (custom request from Perkin Elmer)
- 235 µLFilter Tips, Sterile for Janus (Perkin Elmer, cat. no. 6001289)
- Plate washer: Biotek ELx405 HT
- Centrifuge: Allegra 6 (Beckman Coulter, cat. no. 366802) or PlateFuge (Benchmark Scientific, cat. no. C2000)
- Centrifuge microplate carriers: uPlate Carrier for Rotor SX4750 (Beckman Coulter)
- High content imager; see tables 1-3 for microscopes used by the JUMP Consortium. Our partners also report the Yokogawa CV8000, Yokogawa CQ1, and Perkin Elmer Operetta CLS microscopes have also been used successfully for imaging cell painting experiments. See Jamali et al (in preparation) for more information.
- Plate shaker: Multi-Purpose Rotator (Barnstead Lab-Line, Model 2314)
- Plate reader: EnVision Multilabel Plate Reader (Perkin Elmer, Model 21030010)

**Table 1.** Details of the ImageXpress Micro Confocal channels and stains imaged in the Cell Painting assay.

|  |  |  |  |  |
| --- | --- | --- | --- | --- |
| Dye | Filter (excitation; nm) | Filter (emission; nm) | Organelle or cellular component | CellProfiler channel name |
| Hoechst 33342 | 377/54 | 447/60 | Nucleus | DNA |
| Concanavalin A/Alexa Fluor 488 conjugate | 475/34 | 536/40 | Endoplasmic reticulum | ER |
| SYTO 14 nucleic acid stain | 531/40 | 593/40 | Nucleoli, cytoplasmic RNA | RNA |
| Phalloidin/Alexa Fluor 568 conjugate, wheat-germ agglutinin/Alexa Fluor 555 conjugate | 560/32 | 624/40 | F-actin cytoskeleton, Golgi, plasma membrane | AGP |
| MitoTracker Deep Red | 631/28 | 692/40 | Mitochondria | Mito |

**Table 2.** Details of the PerkinElmer Opera Phenix channels and stains imaged in the Cell Painting assay.

|  |  |  |  |  |
| --- | --- | --- | --- | --- |
| Dye | Filter (excitation; nm) | Filter (emission; nm) | Organelle or cellular component | CellProfiler channel name |
| Hoechst 33342 | 405 | 435-480 | Nucleus | DNA |
| Concanavalin A/Alexa Fluor 488 conjugate | 488 | 500-550 | Endoplasmic reticulum | ER |
| SYTO 14 nucleic acid stain | 488 | 570-630 (widefield) , 515-550 (confocal) | Nucleoli, cytoplasmic RNA | RNA |
| Phalloidin/Alexa Fluor 568 conjugate, wheat-germ agglutinin/Alexa Fluor 555 conjugate | 561 | 570-630 | F-actin cytoskeleton, Golgi, plasma membrane | AGP |
| MitoTracker Deep Red | 640 | 650-760 | Mitochondria | Mito |

**Table 3.** Details of the Yokogawa CV8000 channels and stains imaged in the Cell Painting assay.

| **Dye** | **Filter (excitation; nm)** | **Filter (emission ; nm)** | **Organelle or cellular component** | **Cellprofiler channel name** | **Recommended Channel acquisition order** |
| --- | --- | --- | --- | --- | --- |
| Hoechst 33342 | 405 | 445/45 | Nucleus | DNA | 5 |
| Concanavalin A/Alexa Fluor 488 conjugate | 488 | 525/50 | Endoplasmic reticulum | ER | 4 |
| SYTO 14 nucleic acid stain | 488 | 600/37 | Nucleoli, cytoplasmic RNA | RNA | 3 |
| Phalloidin/Alexa Fluor 568 conjugate, wheat-germ agglutinin/Alexa Fluor 555 conjugate | 561 | 600/37 | F-actin cytoskeleton, Golgi, plasma membrane | AGP | 2 |
| MitoTracker Deep Red | 640 | 676/29 | Mitochondria | Mito | 1 |

### Computational Equipment

- Standard desktop computer. Access to a remote-host computing cluster or cloud-computing platform (optional; recommended if planning to acquire >1,000 fields of view)
- CellProfiler biological image analysis software^20^. Available at http://www.cellprofiler.org
- CellProfiler pipelines: we describe four pipelines in this protocol for illumination correction, segmentation, QC (optional), and feature extraction. The pipelines are available at https://github.com/broadinstitute/imaging-platform-pipelines/tree/master/JUMP_production and were created using CellProfiler 4.1.3. Please see the module notes within the pipelines for Cell Painting-specific documentation. Our Cell Painting wiki (broad.io/CellPaintingWiki) contains a static copy of all files used in the protocol, as well as updates to these files (e.g., to accommodate updated software versions or updated versions of the protocol).
- Raw image data stored in the Cell Painting Gallery on the Registry of Open Data on AWS (https://registry.opendata.aws/cellpainting-gallery/) from chemical perturbations applied to U2OS cells. See download information at https://github.com/carpenterlab/2022_Cimini_NatureProtocols/blob/main/README.md. Note that this plate of images is much larger than we recommend running on a standard desktop computer. To use this data on a local computer we recommend filtering down the images in CellProfiler to a single row, for example. This can be done using the Images module. Replace the rule criterion “Extension Is the extension of an image file” with “File Does Contain r01”, for example.
- Illumination correction images produced by an illumination correction pipeline applied to the U2OS image data. See download information at https://github.com/carpenterlab/2022_Cimini_NatureProtocols/blob/main/README.md.
- A listing of per-cell image features generated by CellProfiler using the analysis pipeline, available at https://github.com/carpenterlab/2022_Cimini_NatureProtocols/blob/main/CellProfiler_features.csv.
- A .csv of per-well profiles generated by running the profiling script, created from the provided sample images. See download information at https://github.com/carpenterlab/2022_Cimini_NatureProtocols/blob/main/README.md.
- A Python^23^ or Conda installation.
- (Optional) KNIME data analytics software^24^ with HCS Tools extension^25^ and sample workflow (See Box 2 for details).
- (Optional) A GUI based SQLite reader, such as DB Browser for SQLite (https://sqlitebrowser.org/)
- (Optional) Plate reader software, such as PerkinElmer EnVision Manager
- (Optional) Data analysis software, such as PRISM from GraphPad Software, LLC

### Reagent Setup (Using PhenoVue Cell Painting JUMP Kit PING 22)

- **PhenoVue 641 Mitochondrial Stain** The product from PerkinElmer (cat. no. PING22) contains 50 µg in each vial. Add 90 µL of DMSO to one vial to make a 1 mM solution. Spin down at 3,200g for 10 sec. Store the solution at -20 °C, protected from light, and use over 3 freeze/thaw cycles.
- **PhenoVue Fluor 555 – WGA** The product from PerkinElmer (cat. no. PING22) contains 0.2 mg in each vial. Add 1.3 ml of dH2O to each to make a 0.15 mg/ml solution. Use a P1000 tip to break up any precipitates in the solution. Store the solution at -20 °C, protected from light, and use it over 3 freeze/thaw cycles.
- **PhenoVue Fluor 488 – Concanavalin A** The product from PerkinElmer (cat. no. PING22) contains 1 mg in each vial. Add 0.5 ml of dH2O to each vial to make a 2 mg/ml solution. Use a P1000 tip to break up any precipitates in the solution. Store the solution at -20 °C, protected from light, and use it over 3 freeze/thaw cycles.
- **PhenoVue Fluor 568 – Phalloidin** The product from PerkinElmer (cat. no. PING22) contains 1 nmol in each vial. Add 150 µL of DMSO to each vial to make a 6.6 µM stock solution. Spin down at 3,200g for 10 sec. Store the solution at -20 °C, protected from light, and use it over 3 freeze/thaw cycles.
- **PhenoVue 512 Nucleic Acid Stain** The product from PerkinElmer (cat. no. PING22) contains 800 nmol/vial. It is a 5 mM solution in DMSO. Spin down at 3,200g for 10 sec. Store the solution at -20 °C, protected from light, and use over 3 freeze/thaw cycles.
- **PhenoVue Hoechst 33342 Nuclear Stain** The product from PerkinElmer (cat. no. PING22) contains 140 µg/vial. It is a 1mg/mL solution in H2O. Spin down at 3,200g for 10 sec. Store the solution at -20 °C, protected from light, and use over 3 freeze/thaw cycles.
- **PhenoVue Dye Diluent A (5x)** The product from PerkinElmer (cat no. PING22) contains 80mL of a 5x stock solution. Dilute 5 times in dH2O to create a 1x HBSS solution with 1% BSA.
- **Triton X-100 solution in HBSS** Add 100 µLof Triton X-100 to 100 ml of HBSS solution to make a 0.1% (vol/vol) Triton X-100 solution. Make fresh solution for each experiment.
- **HBSS (1×)** The product from Invitrogen (cat. no. 14065-056) is 10×. Add 100 ml of HBSS (1×) to 900 ml of water to make HBSS (1×). The 1× solution should preferably be made freshly from the 10× stock solution, but it can also be stored at 4 °C for up to 1 week. If storing, filter the HBSS (1×) with a 0.22-?m filter
- **Live-cell Mitochondrial staining solution** Prepare Mitochondrial staining solution by adding 150 µLof PhenoVue 641 Mitochondrial 1mM stock solution to 100 ml of prewarmed cell culture medium (preferred method) or HBSS for a working concentration of 1.5 µM to end with a final staining concentration of 500 nM. Make fresh solution for each staining session. We recommend not increasing the concentration of the staining solution any further, to avoid small changes in dispensing volume leading to high variability in final effective concentration. Keep the solution wrapped in foil and away from light.
- **Phalloidin, concanavalin A, Hoechst, WGA, and Nucleic Acid Stain 512 staining solution** To make 100 ml of stain solution, add 125 µL of 6.6 µM phalloidin stock solution, 250 µL of 2 mg/mL concanavalin A stock solution, 100 µLof 1 mg/mL Hoechst stock solution, 1 mL of 0.15 mg/mL WGA stock solution, and 120 µLof 5mM Nucleic Acid Stain 512 stock solution in 1x HBSS (1% BSA and 0.1% Triton X-100). Keep the solution wrapped in foil and away from light.
- **Fixation Solution** To create a 16% solution of PFA, dilute 32% PFA from Electron Microscopy Sciences (cat no. 15740-S) 1:1 with distilled water. Handle in fume hood.
- **Compound library** Dissolve the compounds in DMSO to yield the desired molarity. Do not exceed a final DMSO concentration of 0.5% in the destination well. Seal and store it at -20 °C for long-term storage or at room temperature (RT; 21 °C) for up to 6 months in a dessicator; other common compound management solutions may also be used45.

### REAGENT SETUP NEEDED FOR LENTIVIRAL TRANSDUCTION

- **Growth media** To 500 mL of DMEM or McCoy’s base media add 50 mL FBS and 5 mL pen/strep
- **Polybrene solution** Prepare growth media containing 16 ug/mL polybrene from a polybrene stock of 8 mg/mL based on the total volume needed to seed 10 µLper 384w. Different cell types will require varying concentrations of polybrene for aid in lentiviral transduction and varying concentrations should be assessed for cell toxicity, see Supplemental Materials.
- **Blasticidin solution** For use with lentiviral expression vectors containing a blasticidin resistance cassette. Prepare cell culture media containing a final concentration of 16 ug/mL blasticidin from a blasticidin stock of 10 mg/mL based on the total volume needed to seed 50 µLper 384w. Different cell types will require varying concentrations of blasticidin for complete antibiotic selection, see Supplemental Materials.
- **Puromycin solution** For use with lentiviral expression vectors containing a puromycin resistance cassette. Prepare cell culture media containing a final concentration of 0.75 ug/mL puromycin from a puromycin stock of 10 mg/mL based on the total volume needed to seed 50 µLper 384w. Different cell types will require varying concentrations of puromycin for complete antibiotic selection, see Supplemental Materials.
- **Control compound spike-ins** Initially prepare 10 mM TC-S 7004 stock in DMSO from powder. Aliquot 5 µLof 10 mM stocks of compounds into PCR strip tubes for single use to avoid freeze/thaw cycles, then store AMG-900, LY2109761, and TC-S 7004 at -20C and Quinidine at -80C. For manual spike-in to ORF-treated 384w plates, add 1 µLof 10 mM DMSO stock + 1 µL8 mg/mL polybrene to 498 µLcell culture media and mix, final concentration 20 uM. Then add 10 µLof 20 uM stock to each 384w seeded at 30 uL/well for a final concentration of 5 uM.
- **(Optional) CellTiter-Glo** Bring to room temperature the CellTiter-Glo reagents, including the CellTiter-Glo Substrate and CellTiter-Glo Buffer. Once reagents have come to room temperature, aspirate the total volume of substrate present in the tube, either 10, 100, or 500 mL depending on the size kit that was purchased. Dispense that volume into the container containing the powered buffer. Mix by gently pipetting up and down until the powder has thoroughly dissolved. It is recommended to then cover the container or transfer to another container covered in foil, due to the reagents being light-sensitive.

### Equipment Setup

- **Microscope selectio** The Cell Painting assay has been applied using both wide-field and confocal microscopy. Confocal microscopes are able to achieve higher image contrast (and hence increased cellular feature definition and improved object segmentation) by rejecting light originating from out-of-focus planes of field. However, as compared with wide-field microscopes, confocal microscopes possess a limited number of excitation wavelengths available for use, have typically higher purchase prices, which may be prohibitive for smaller research groups, and are traditionally of lower throughput.

We have used three High Content Imaging Systems: an ImageXpress Micro XLS (Molecular Devices), an Opera Phenix (PerkinElmer), and Yokogawa **CV8000**. The images are captured in five fluorescent channels given in Table 1. Note that while the Opera Phenix is capable of imaging in both wide-field and confocal modes, in confocal mode it cannot adjust its emission filter such that SYTO14 can be captured separately from Alexa 488.

If multiple microscopes must be used, we recommend imaging one full replicate all on one microscope, as opposed to arbitrarily assigning plates to different instruments as the experiment proceeds. The rationale is to avoid imager-induced batch effects. If the differences between perturbations are marked, then post-acquisition normalization will probably be effective (see ‘Normalization of morphological features across plates’ in the PROCEDURE section for more details).

However, if the morphological effects to be measured are subtle, normalization may not be sufficient, and the similarities in the collected image features will more likely reflect the different image acquisition than the underlying biological perturbations.

**CRITICAL** The same microscope should be used for imaging all microtiter plates during an experiment. We do not recommend switching microscopes midstream because lamp intensities, filter patterns, and other subtleties can be quite different even between supposedly identical microscope setups.

- **Automated image acquisition setting** Each channel should be captured as an individual grayscale image. No further pre-processing should be performed on the images before analysis.

The choice of objective magnification is important as there is a trade-off between increased image feature resolution at higher magnifications (therefore enabling more specific quantification of certain organelles) versus a smaller field of view and hence fewer cells imaged (therefore decreasing throughput and statistical power for profile generation). Acquiring more fields of view can mitigate the latter consideration, but at the cost of a substantial increase in image acquisition and computational processing time, especially for those who do not have access to computing cluster resources. We have found that using a 20× water-immersion objective (NA 1.0) sufficiently balances all competing issues.

Typically, nine sites are collected per well in a 3 × 3 site layout, at 20× magnification and 2× binning. If time permits, more sites can be imaged in order to increase well coverage and to improve sample statistics; it is best to capture as many cells as possible.

If possible on your microscope, adjusting the relative acquisition heights of each channel may lead to optimal capture of each channel’s relevant cell structures. The JUMP ORF production data had 2µm total Z difference between channels captured at the lowest Z position and channels captured at the highest Z position.

The order in which the channels are imaged may have an impact on the likelihood of photobleaching during the experiment; photobleaching manifests as a decay in the fluorescence signal intensity over time with repeated illumination. As the emission wavelengths for the chosen fluorophores are broad and in close proximity to each other, photobleaching may occur for the low-intensity dyes as they are irradiated by the lower-wavelength light. To mitigate this effect, we recommend imaging the five channels in order of decreasing excitation wavelength. For Opera Phenix instruments equipped with more than one camera, we recommend to separate all channels.

**CRITICAL** Be sure that the images are not saturated. Generally, set exposure times such that a typical image uses roughly 50% of the dynamic range. For example, because the pixel intensities will range from 0 to 65,535 for a 16-bit image, a rule of thumb is for the typical sample to yield a maximum intensity of ∼32,000. This guideline will prevent saturation (i.e., reaching the value 65,535) from samples that are brighter because of a perturbation.

Do not use shading correction if you are using the recommended CellProfiler workflow for image analysis, as the background illumination heterogeneities will be corrected post-acquisition using the CellProfiler software.

Before beginning the complete imaging run, it is useful to capture images from three to five wells at a few different locations across the plate, in order to confirm that the microscope is operating as expected and the acquisition settings are optimal for the experiment and cell line at hand. See the Computational Equipment section for a link to an example image data set.

We recommend including the acquisition of at least one brightfield channel, which can be used in feature extraction but may also be useful for downstream applications such as fluorescence channel prediction. The JUMP ORF production captured 3 Z positions - one equal to the lowest fluorescence position (Brightfield), one 5µm above that position (BFHigh), and one 5µm below that position (BFLow).

**CRITICAL** Avoid capturing the edges of the well in the images, particularly if a large number of sites per well are imaged. Although it is feasible to remove the well edges from the images post-acquisition using image processing approaches, such methods are challenging and best avoided.

- **Image processing software** CellProfiler biological image analysis software is used to extract per-cell morphology feature data from the Cell Painting images, and can optionally be used to extract per-image QC metrics (see Box 2). The software and associated pipelines are designed to handle both low- and high-throughput analysis, but we routinely run this software as part of this protocol on thousands, even millions, of imaged fields of view.

To download and install the open-source CellProfiler software, go to http://www.cellprofiler.org, follow the download links for CellProfiler, and follow the installation instructions. The current version at the time of writing is 4.2.1.

This protocol assumes basic knowledge of the CellProfiler image analysis software package. Extensive online documentation and tutorials can be found at http://www.cellprofiler.org/. In addition, the ‘?’ buttons within CellProfiler’s interface provide detailed help. The pipelines used here are compatible with CellProfiler version 4.1.3 and above.

This protocol uses three CellProfiler pipelines to perform the following tasks: illumination correction, segmentation, and morphological feature extraction. Additionally, you can use an optional QC pipeline to quantify image quality. See the Computational Equipment section for a link to the CellProfiler pipelines used in this protocol.

Each module of the pipelines is annotated with details about the purpose of the module and considerations in making adjustments to the settings. The annotations may be found at the top of the settings, in the panel labeled ‘Module notes’.

The pipelines are configured on the assumption that the image files follow the nomenclature of the Perkin Elmer Opera Phenix system, in which the plate/well/site metadata are encoded as part of the filename. The plate and well metadata in particular are essential because CellProfiler uses the plate metadata in order to process the images on a per-plate basis, and the plate and well metadata are needed for linking the plate layout information with the images for the downstream profiling analysis. Therefore, images coming from a different acquisition system may require adjustments to the Metadata module to capture this information; please refer to the help function for this module for more details or our video tutorial walkthrough of the Input modules https://youtu.be/Z_pUWuOV06Q.

The QC and morphological feature extraction pipelines are set to write out cellular features to a .csv for each well and site, respectively, using the ExportToSpreadsheet module. We provide Python scripts to generate per-well profiles from the extracted features.

- **Computing system** Small batches of images can be processed on most modern desktop computers. If the number of images to analyze is sufficiently large (e.g., >∼1,000 images), processing time on a single computer becomes resource-limiting. For large batches of images, we recommend using a computing cluster if available, such as a high-performance server farm, or a cloud-computing platform such as Amazon Web Services (AWS). Substantial setup effort is required for both cluster computing and cloud-distributed computing, and will probably require enlisting the help of your IT department. Please refer to our GitHub webpage (https://github.com/CellProfiler/CellProfiler/wiki/Adapting-CellProfiler-to-a-LIMS-environment) for more details on cluster computing, and our Distributed CellProfiler GitHub repository documentation (https://distributedscience.github.io/Distributed-CellProfiler/overview.html) for more details on cloud computing using AWS.

## Procedure

### Cell culture

**TIMING** variable; 2–7 d

**CRITICAL** The following cell-plating procedure has been validated for many cell types; each step may need adjustment depending on local conditions or alternative cell types. We have included recommended optional steps for experiments involving small-molecule library treatment, ORF overexpression, and CRISPR knockdown.

**CRITICAL** Check the wiki at GitHub for any updates to the Cell Painting protocol: broad.io/CellPaintingWiki

Prepare cells for seeding according to known best practices for the cell type of choice. For most high-content applications, a black plate with a clear, flat bottom for cell culture is appropriate. The following protocol has been validated for use on U2OS cells in Corning 384-well 200-nm-thick glass-bottom plates. (Optional) White plates with a clear, flat bottom for cell culture are also appropriate for any cell viability assay using CellTiter-Glo.

1. Grow cells to near confluence (∼80%) in a T-175 tissue culture flask.
2. (Optional) If you are performing experiments that involve the addition of compounds (Step 8A), prepare the compound library according to the instructions in the Reagent Setup.
3. Rinse the cells with PBS without Ca2+ or Mg2+.
4. Add 3 ml of Trypsin-EDTA solution and incubate the cells at 37 °C until the cells have detached. This should occur within 3–5 min.
5. Add 4 ml of growth medium to deactivate the trypsin, and collect the cell suspension into a conical tube.
6. Wash the tissue culture flask with an additional 5 ml growth media, add the wash to the same conical tube with the cell suspension, and mix thoroughly but gently.
7. Determine the live-cell concentration using standard methods (hemocytometer or cell counter).
8. Dilute U2OS cells to 50,000 live cells per ml in media, and dispense 30 µL (2,000 live U2OS cells) into each well of the 384-well plates. For large-scale Cell Painting assays, we recommend the use of an automated liquid-handling system. Different cell types and growth conditions will require variations in seeding density; typical ranges will vary from 1,500 to 3,000 cells per well. (Optional) Dispense cells into white plates if any assays are to be performed with CellTiter-Glo.
9. Keep cell plates at RT for about 1h before proceeding with the small molecule, viral ORF overexpression or viral CRISPR knockdown or transfer them to the incubator. **CRITICAL STEP** Adequately resuspend the cell mixture to ensure a homogeneous cell suspension before each dispensation. It is not uncommon for cells to rapidly settle in their reservoir, resulting in plate-to-plate variation in cell numbers. If you are using a liquid handler with a multi-dispense function, be sure to adequately sterilize and prime the dispensing cassette and/or dispense at least 10 µL of cell suspension back into the reservoir before dispensing the cells into culture plates; the latter is helpful if cells or reagents are sticking to the tubing. **CRITICAL STEP** When handling liquid for many plates with one set of tips, confirm that no residual bubbles within the tips touch the head of the liquid handler during aspiration, in order to ensure accurate liquid dispensation.

### Treatment with a small-molecule library, viral ORF overexpression library, or viral CRISPR knockdown library

10. If you are performing treatments with a small-molecule library, please follow option A. If you are performing viral CRISPR knockdown, please follow option B. If you are performing viral ORF overexpression, please follow option C. For instructions on using siRNA transfection, please refer to the 2016 protocol. To facilitate alignment of data across batches, no matter which modality you use for your own data we recommend in each batch of Cell Painting plates you prepare both an additional negative control plate (using only DMSO as the treatment for cells) and an additional positive control plate (we recommend a control compound plate such as the JUMP-Target-1 or JUMP-Target-2 plates).
  a. Addition of a small-molecule library **TIMING** variable; ∼2–3 d for one batch experiment of 384-well plates
    i. Allow plates to sit on a flat, level surface at RT for 1–2 h after seeding to reduce plate edge effects46.
    ii. Put the plates into the incubator (37 °C, 5% CO2, 90–95% humidity). To reduce plate edge effects produced by incubator temperature variations and media evaporation, we recommend either spacing out the plates in the incubator or using racks with ‘dummy’ plates filled with liquid placed on the top and bottom. We also recommend rotating the plates/stacks within the incubator to avoid positional effects.
    iii. Add compounds to cells using a pin tool or other liquid handler (e.g. the Agilent Bravo automated pipetting platform or the Beckman Coulter Echo acoustic liquid handling system). Compounds may be added either 24 or 48 h before staining and fixation, but this timing should be adjusted depending upon the growth rate of the cell types being used and the biological processes under consideration. Recursion Pharmaceuticals typically adds compounds to cells in an environment that is antibiotic-free (to avoid perturbations arising from complex antibiotic–drug interactions) and low-serum (to synchronize cell state). To ensure adequate mixture of compounds in solution, we recommend that compounds be mixed well in the culture medium before adding them to the cells. **CRITICAL STEP** Plates should be stacked in the incubator no more than three-high with PBS 384-well plates on top and bottom and rotated every 24 hours to help prevent edge effects. **CRITICAL STEP** Ensure that the liquid handler aspiration and dispense speeds are set to the slowest possible setting to avoid distributing the cells after the initial seeding.
  b. Addition of viral CRISPR knockdown constructs **TIMING** variable; ∼4–6 d for one batch experiment of 384-well plates
    i. Allow plates to sit on a flat, level surface at RT for 1–2 h after seeding to reduce plate edge effects.
    ii. CRISPR knockdown lentivirus is pre-prepared in 384-well plates at -80C. Let thaw on wet ice, then quickly pulse spin plate to prevent any virus from clinging to plate seal. Place back on ice until needed.
    iii. Using a liquid handler, add 10 uL/384-well of growth media containing polybrene as described in the Reagent Setup.
    iv. Immediately after polybrene addition, add the appropriate volume of CRISPR knockdown lentivirus per 384-well using a liquid handler. Different cell types and lentiviral expression vectors will require variations in lentivirus volume; typical ranges will vary from 0.5 to 4 µLper well.
    v. Gently tap the plates to ensure an even distribution of cells across each well.
    vi. After lentiviral transduction, centrifuge the 384-well plate(s) for 30 minutes at 1,178 g at 37°C.
    vii. After centrifugation, place the cell plates in an incubator (37°C, 5% CO2).
    viii. 24 hours post-lentiviral transduction, remove 50 µLof media and replace with 50 µLfresh growth media and return to the incubator. (Optional) If assessing the efficiency of the lentiviral transduction as a quality control measure, for two white cell plates, replace the growth media in one with 50 µLmedia and the other with 50 µLmedia containing puromycin as described in the Reagent Setup.
    ix. The timing post viral CRISPR knockdown transduction and prior to downstream assays may either be 96 or 144 h, but this timing should be adjusted depending upon the growth rate of the cell types being used and the biological processes under consideration.
    x. If performing downstream assays after 144 h: At 96 h post-viral CRISPR knockdown transduction, remove 50 µLof media and replace with 50 µLfresh growth media or (optional) 50 µLmedia containing puromycin to a white cell plate and return to the incubator.
    xi. After the predetermined number of hours, proceed with the appropriate subsequent steps for cell fixation, staining and imaging or (optional) determining the lentiviral transduction efficiency.
    xii. (Optional) The transduction efficiency is determined by adding 10 µLper 384-well room temperature CellTiter-Glo to two white cells plates, one with puromycin treatment and one without puromycin treatment. Cover the plates with aluminum foil and put on a shaker at low speed for 15 minutes. Then image the plates using a standard plate reader such as Envision Multilabel Reader (Perkin Elmer). **CRITICAL STEP** Plates should be stacked in the incubator no more than three-high with PBS 384-well plates on top and bottom and rotated every 24 hours to help prevent edge effects. **CRITICAL STEP** Ensure that the liquid handler aspiration and dispense speeds are set to the slowest possible setting to avoid distributing the cells after the initial seeding.
  c. Addition of viral overexpression constructs **TIMING** variable; ∼2–4 d for one batch experiment of 384-well plates
    i. Allow plates to sit on a flat, level surface at RT for 1–2 h after seeding to reduce plate edge effects.
    ii. ORF overexpression lentivirus is pre-prepared in 384-well plates at -80C. Let thaw on wet ice, then quickly pulse spin plate to prevent any virus from clinging to plate seal. Place back on ice until needed.
    iii. Using a liquid handler, add 10 uL/384-well of growth media containing polybrene as described in the Reagent Setup.
    iv. Immediately after polybrene addition, add the appropriate volume of ORF overexpression lentivirus per 384-well using a liquid handler. Different cell types and lentiviral expression vectors will require variations in lentivirus volume; typical ranges will vary from 0.5 to 4 µLper well.
    v. (Optional) Control compounds can be spiked into well(s) that do not contain ORF overexpression perturbations if desired. See the Reagent Setup.
    vi. Gently tap the plates to ensure an even distribution of cells across each well.
    vii. After lentiviral transduction, centrifuge the 384-well plate(s) for 30 minutes at 1,178 g at 37°C.
    viii. After centrifugation, place the cell plates in an incubator (37°C, 5% CO2).
    ix. 24 hours post-lentiviral transduction, remove 40 µLof media and replace with 40 µLfresh growth media and return to the incubator. (Optional) If assessing the efficiency of the lentiviral transduction as a quality control measure, for two white cell plates, replace the growth media in one with 50 µLmedia and the other with 50 µLmedia containing blasticidin as described in the Reagent Setup.
    x. The timing post viral ORF overexpression transduction and prior to downstream assays may either be 48 or 96 h, but this timing should be adjusted depending upon the growth rate of the cell types being used and the biological processes under consideration. After the predetermined number of hours, proceed with the appropriate subsequent steps for cell fixation, staining and imaging or (optional) determining the lentiviral transduction efficiency.
    xi. (Optional) The transduction efficiency is determined by adding 10 µLper 384-well room temperature CellTiter-Glo to two white cells plates, one with blasticidin treatment and one without blasticidin treatment. Cover the plates with aluminum foil and put on a shaker at low speed for 15 minutes. Then image the plates using a standard plate reader such as Envision Multilabel Reader (Perkin Elmer). **CRITICAL STEP** Plates should be stacked in the incubator no more than three-high with PBS 384-well plates on top and bottom and rotated every 24 hours to help prevent edge effects. **CRITICAL STEP** Ensure that the liquid handler aspiration and dispense speeds are set to the slowest possible setting to avoid distributing the cells after the initial seeding.

### Staining and fixation

**TIMING** variable; 2.5–3 h for one batch experiment of 384-well plates

11. Prepare the following for all plates in advance of initiating the staining process:
  a. The live-cell PhenoVue 641 Mitochondrial staining solution in cell media or HBSS
  b. The fixation solution containing 16% w/v PFA in distilled water
  c. The staining & permeabilizing solution containing PerkinElmer PhenoVue dyes Hoechst 33342, Fluor 488 Concanavalin A, 512 Nucleic Acid Stain, Fluor 555 WGA, and Fluor 568 Phalloidin in 1X PhenoVue Dye Diluent A with 0.1% Triton
12. Leaving growth media in place to minimize disturbance to the live cells, add 20µLof the mitochondrial staining solution over the top of each well to a final well volume of 60uL.
  a. Place a stir bar in the source bottle to prevent dye from settling and causing adverse plate patterns.
  b. We recommend using a metal tip cassette to prevent bubbling and allow for a more even dispense of dye. **TIP:** if using a liquid handler such as a Combi multidrop, be aware that dye can settle in tubing if there is a significant lapse of time between dispenses (>2min/plate). This may cause adverse plate patterns. We recommend introducing a repriming step to clear the dyes from the tube and introduce fresh dye from the source bottle. **TIP:** If plate patterns persist, the addition of an initial blank plate (no cells present) may alleviate the intensity of patterns.
13. If necessary, centrifuge the plate (500g at RT for 1 min) after adding stain solutions to ensure that there are no bubbles in the bottoms of the wells.
14. Incubate the plates for 30 min in the dark at 37 °C. **TIP:** Note, once mitochondrial staining solution is added, keep plates in dark for the rest of the experiment.
15. To fix the cells, add 20µLof 16% (w/v) methanol-free PFA on top of each well to bring each well to a final volume of 80µLwith final concentration of 4% PFA (v/v).
16. Incubate the plates in the dark at RT for 20 min.
17. Wash the plates four times with 80 µLof 1x HBSS. Include a final aspiration step.
18. To each well, add 20 µL of the staining & permeabilizing solution.
  a. Place a stir bar in the source bottle to prevent dye from settling and causing adverse plate patterns.
  b. If experiencing bubbling, we recommend using a metal tipped cassette.
19. Incubate the plates in the dark at RT for 30 min.
20. Wash the plates four times with 80µLof 1x HBSS. Leave a final volume of 80µLof 1x HBSS in each well.
21. Seal the plates with adhesive foil and store them at 4°C in the dark. **TIP:** If plates are to be stored long-term, 0.05% sodium azide can be added to mitigate contamination. **CAUTION:** Sodium azide may cause damage to organs through prolonged or repeated exposure and is fatal if swallowed, in contact with skin or if inhaled.

### Automated image acquisition

**TIMING** variable; 1–3.5 h per 384-well plate

22. Acquire images from the microtiter plates using the high content imager. For large-scale Cell Painting assays, we recommend the use of an automated microplate handling system.
23. Set up the microscope acquisition settings as described in the Equipment Setup.
24. Start the automated imaging sequence according to the microscope manufacturer’s instructions. SEE ALSO TROUBLESHOOTING

### Morphological image feature extraction from microscopy data

**TIMING** variable; 20 h per batch of 384-well plates

**Perform illumination correction** to improve fluorescence intensity measurements:

25. Start CellProfiler.
26. Load the illumination correction pipeline into CellProfiler by selecting File > Import > Pipeline from File from the CellProfiler main menu and then selecting illumination.cppipe or dragging and dropping the pipeline from your file browser into the pipeline panel on the left of the interface. **CRITICAL STEP** Nonhomogeneous illumination introduced by microscopy optics can result in errors in cellular feature identification and can degrade the accuracy of intensity-based measurements. This is an especially important problem in light of the subtle phenotypic signatures that image-based profiling aims to capture. Nonhomogeneous illumination can occur even when fiber-optic light sources are used and even if the automated microscope is set up to perform illumination correction. The use of a uniformly fluorescent reference image (‘white-referencing’), although common, is not suitable for high-throughput screening. A retrospective method to correct all acquired images on a per-channel, per-plate basis is therefore recommended^26^; the illumination pipeline takes this approach.
27. Select the Images input module in the ‘Input modules’ panel to the top-left of the interface. From your file browser, drag and drop the folder(s) containing your raw images into the ‘File list’ panel. See the Computational Equipment section for a link to raw image files that can be used as an example in this protocol. Note that the
28. Click the ‘Output settings’ button at the bottom-left of the interface. Select an appropriate ‘Default Output Folder’ into which the illumination correction images will be saved.
29. Save the current settings to a project (.cpproj) file containing the pipeline, the list of input images, and the output location by selecting File > Save Project. Enter the desired project filename in the dialog box that appears. This is not necessary for the running of this step, however it functions as a snapshot of your complete setup, allowing you to directly replicate your work by loading this .cpproj instead of the .cppipe you started with.
30. Press the ‘Analyze Images’ button at the bottom-left of the interface. A progress bar in the bottom-right will indicate the estimated time of completion. The end result of this step will be a collection of illumination correction images in the Default Output Folder, one for each plate and channel. We have provided an example set of images for comparison on our Cell Painting wiki (see Computational Equipment for details). **CRITICAL STEP** This step assumes that you will be running the illumination correction pipeline locally on your computer. If your institution has a shared high-performance computing cluster, such as a high-performance server farm, or a cloud-computing platform such as Amazon Web Services (AWS), we recommend executing the pipeline on the cluster as a batch process—i.e., a series of smaller processes entered at the command line; this will result in much more efficient processing. Enlist the help of your institution’s IT department to find out whether this is an option and what resources are available. If choosing this option, carry out the instructions in Box 1, describing modifications to the pipeline to run it as a batch process. **Configure the segmentation pipeline** to optimize cell and nuclei segmentation:
31. Start CellProfiler, if you are not already running it.
32. Load the Segmentation pipeline into CellProfiler by selecting File > Import > Pipeline from File from the CellProfiler main menu and selecting segmentation.cppipe or dragging and dropping the pipeline from your file browser into the pipeline panel on the left of the interface. **CRITICAL STEP** Because capturing subtle phenotypes is important for profiling, accurate nuclei and cell body identification is essential for success. This pipeline enables troubleshooting of nuclei and cell segmentation across an entire batch by outputting an image per well with nuclei and cell objects overlaid for visual inspection. The optimal parameters determined in this pipeline must be manually transferred to the analysis pipeline.
33. Select the Images input module in the ‘Input modules’ panel to the top-left of the interface. From your file browser, drag and drop the folder(s) containing your raw images into the ‘File list’ panel.
34. Enter CellProfiler’s Test mode using the Start Test Mode button. Examine the outputs of IdentifyPrimaryObjects and IdentifySecondaryObjects for a few images to make sure that the boundaries generally match expectations. Under the ‘Test’ menu item, there are options for selecting sites for examination. We recommend either randomly sampling images for inspection (via ‘Random Image set’) and/or selecting specific sites (via ‘Choose Image Set’) from both negative control wells and specific treatment locations from the plates. The rationale is to check a wide variety of treatment-induced phenotypes to ensure that the pipeline will generate accurate results. The CellProfiler website contains resources and tutorials on how to optimize an image analysis pipeline. See also Table 4. Troubleshooting.

**Table 4.** Troubleshooting table.
35. Press the ‘Analyze Images’ button at the bottom-left of the interface. A progress bar in the bottom-right will indicate the estimated time of completion. The end result of this step will be a collection of images with nuclei and cell segmentations overlaid on them in a ‘Segmentations’ folder in the Default Output Folder. We have provided an example set of images for comparison on our Cell Painting wiki (see Computational Equipment for details). **CRITICAL STEP** This step assumes that you will be running the segmentation pipeline locally on your computer. If your institution has a shared high-performance computing cluster, such as a high-performance server farm, or a cloud-computing platform such as Amazon Web Services (AWS), we recommend executing the pipeline on the cluster as a batch process—i.e., a series of smaller processes entered at the command line; this will result in much more efficient processing. Enlist the help of your institution’s IT department to find out whether this is an option and what resources are available. If choosing this option, carry out the instructions in Box 1, describing modifications to the pipeline to run it as a batch process.
36. Visually inspect your output segmentation images. Open the segmented images in your preferred image viewer (we recommend Fiji^27^/ImageJ^28^) and decide if the computationally determined Nuclei and Cell objects correspond to what you see by your eye across your batch. Perfection is nigh impossible, but you should agree with the vast majority of segmentations. A rule of thumb is that you should see roughly the same number of objects over-segmented (one object labeled as two or more) as under-segmented (two or more objects labeled as one). If not satisfied with the segmentation, repeat the segmentation pipeline workflow with different segmentation parameters. To simplify image visualization, particularly in larger datasets, you may wish to create whole-plate montages of your images using methods described in Dobson et al ^17^; images with annotations can be saved as SVG files via a Fiji plugin^29^. The ANTICIPATED RESULTS section outlines the expected nuclei and cell identification quality. **Run quality control** to extract image quality measurements: See Box 2 for this optional step. **Run the final analysis pipeline** to extract morphological features:
37. Start CellProfiler, if you are not already running it.
38. Load the analysis pipeline into CellProfiler by selecting File > Import > Pipeline from File from the CellProfiler main menu and selecting analysis.cppipe or dragging and dropping the pipeline from your file browser into the pipeline panel on the left of the interface.
39. Select the Images input module in the ‘Input modules’ panel to the top-left of the interface. From your file browser, drag and drop the folder(s) containing your raw images into the ‘File list’ panel.
40. If you needed to change segmentation parameters in your Segmentation pipeline for accurate object identification, make those same changes to the IdentifyPrimaryObjects and IdentifySecondaryObjects modules for the identification of NucleiIncludingEdges and CellsIncludingEdges, respectively.
41. If you did not run a separate QC pipeline and would like to measure image-level quality metrics, check the two MeasureImageQuality modules at the beginning of the pipeline to include those measurements in the analysis pipeline.
42. Click the ‘Output settings’ button at the bottom-left of the interface. Select an appropriate ‘Default Output Folder’ into which the analysis data will be saved.
43. Save the current settings to a project (.cpproj) file containing the pipeline, the list of input images, and the output location by selecting File > Save Project. Enter the desired project filename in the dialog box that appears. This is not necessary for the running of this step, however it functions as a snapshot of your complete setup, allowing you to directly replicate your work by loading this .cpproj instead of the .cppipe you started with.
44. **CRITICAL STEP** Press the ‘Analyze Images’ button at the bottom-left of the interface. A progress bar in the bottom-right will indicate the estimated time of completion. The pipeline will identify the nuclei from the Hoechst-stained image (referred to as ‘DNA’ in CellProfiler), then it will use the nuclei to guide identification of the cell boundaries using the SYTO 14–stained image (‘RNA’ in CellProfiler), and then it will use both of these features to identify the cytoplasm. The pipeline then measures the morphology, intensity, texture, and adjacency statistics of the nuclei, cell body, and cytoplasm, and it outputs the results to a series of .csvs. See the Equipment section for a link to a listing of the image features measured for each cell. **CRITICAL STEP** This step assumes that you will be running the image analysis pipeline locally on your computer, which generally is recommended only for experiments with <1,000 fields of view. If your institution has a shared high-performance computing cluster, such as a high-performance server farm, or a cloud-computing platform such as Amazon Web Services (AWS), we recommend executing the pipeline on the cluster as a batch process—i.e., a series of smaller processes entered at the command line; this will result in much more efficient processing. Enlist the help of your institution’s IT department to find out whether this is an option and what resources are available. If choosing this option, carry out the instructions in Box 1, which describes modifications to the pipeline to run it as a batch process.

| Step | Problem | Possible reason | Solution |
| --- | --- | --- | --- |
| 8 | Cells are not evenly plated across all wells | Air bubbles during cell dispensing, cell solutions settling after suspension but before seeding | Make sure the cell suspension is not settling or clumping by using gentle stirring or agitation, ideally via magnetic stir plate; prime the seeding cassettes; adjust pressure and dispense speed of the cell suspension; if needed, change seeding cassettes. |
| 20 | Plated cells are lost unevenly across the plate | Problems with the automated plate washer. | Check plate washer's manifold alignment and aspiration speed and height; check also for clogged needles. |
|  |  | Problems with the reagent dispenser. | Check height and speed of dispensation; use angled reagent dispensing where possible |
| 24 | The images contain bright, slender, or punctate artifacts that appear in multiple wells, across multiple channels. Too many of these artifacts can adversely affect nuclei and cell body identification and measurement | The washing reagents are contaminated with fibers—e.g., from clothing or dust | Filter the washing solutions and diluents before use. Prepare plates in a clean, dust-minimal environment, and dust the bottom of plates before imaging.<br>While ideally fixed before acquisition, if your images do have a significant amount of debris, you may wish to add into your pipelines additional IdentifyPrimaryObjects module/s to detect the debris and then mask the debris from your images using MaskImage modules before identifying your nuclei and cell objects. |
|  |  | Bacterial contamination of the plates | Check the plate washer and other automation tools to find the source of contamination and clean per manufacturer instructions. Image plates soon after fixing and staining to avoid long-term growth of contaminants. |
| 24 | Staining intensities are uneven across a multiwell plate. | Artifacts in dispensing dyes. | Careful preparation of stock solutions to ensure homogeneity. Prime automated dispensers with staining solution. Check dispensers for clogs. |
|  |  | Autofluorescent compounds | Adjust washing procedure to minimize residual volume while maintaining attached cells. In case of insufficient washing, additional washing steps might be required. |
| 24 | High background fluorescence | Inadequate performance of the cell washer | Adjust washing procedure to minimize residual volume while maintaining attached cells. In case of insufficient washing, additional washing steps might be required. |
|  |  | Autofluorescent compounds | Adjust washing procedure to minimize residual volume while maintaining attached cells. In case of insufficient washing, additional washing steps might be required. |
| 24 | Bad image quality for the whole plate or parts of the plate | Plate manufacturing errors | Ensure that autofocus is working well on the imagers to avoid artifacts due to drifts in plate bottom thickness. Use Z-projection to prevent well-to-well drifts. When possible, buy plates from a single lot to avoid batch-to-batch quality issues. |
|  |  | Imager astigmatism/optical alignment | Regular profiling of the imaging system should be performed using fluorescent beads; the system should be recalibrated as needed per manufacturer instructions. Special care should be taken on systems with multiple cameras. Acquiring multiple z-planes can help to counteract focal artifacts due to issues with optical alignment or sample tilt/flatness. Ensuring that FOVs are close to the center of each well in the microplate can also help to minimize the variations in the bottom surface of the microplate well. |
|  |  | Debris on objectives/in light path | Perform preventative maintenance and conduct visual inspection of sample images before running larger scale campaigns and cleaning according to manufacturers recommendations |
| 34 | The identified nuclei or cell bodies do not reflect the actual boundaries of the stained nuclei or cells in the image | The settings in the IdentifyPrimaryObjects or IdentifySecondaryObjects modules (for nuclei and cell identification, respectively) were optimized for U2OS cells imaged on a particular microscope at a particular magnification, and may be inappropriate for different experimental conditions | Cell lines with different morphological features may require additional optimization of the pipeline identification modules. After launching CellProfiler and loading the feature extraction pipeline, see the Module Notes in the main window of CellProfiler for more details on relevant settings for each module. Visual inspection is needed to confirm that the settings conform to expected results. If you encounter difficulties in adjusting the pipeline settings for this task, we recommend consulting the moderated forum at http://forum.image.sc/ for assistance |
|  |  | Cell confluency can have a profound impact on your ability to determine cell boundaries and thus segment cells. Segmentation algorithms can struggle if there is no background (i.e. blank space) in an image and particularly confluent cells may grow on top of each other making segmentation impossible. | Experiment with plating your cells at different concentrations to find an appropriate balance between maximizing data and leaving some background in your images while avoiding cells growing on top of each other.<br>While ideally fixed before acquisition, if your images do have heavily confluent regions preventing accurate segmentation, you may wish to add into your pipelines an additional IdentifyPrimaryObjects module to detect the confluent region and then mask the confluent region from your images using MaskImage modules before identifying your nuclei and cell objects. |

### Creation, normalization, and feature reduction of per-well profiles

All steps in this section are covered in more detail and are continuously updated in the profiling-handbook^30^

**TIMING** 8-16 h per 384-well plate

45. Set up a terminal for bash scripting as well as a language you are comfortable writing scientific code in; most of our tooling is in Python^23^ since it has a large number of useful packages you can use to write ^31–37^ or execute^38^ code per our templates but it is not mandatory.
46. Create per-well mean aggregated profiles using the pycytominer^39^ command ’collate.py’ If you have run your CellProfiler pipeline in a cluster computing environment, you can also use this same script to aggregate multiple CSV files into a single SQLite database before aggregation using the cytominer-database^40^ package. This step can take 8-16 hours per plate per plate to run; if running multiple plates, it is strongly recommended to use parallel^41^ or a similar tool to parallelize profile creation.
47. Create a metadata file describing the treatments on each plate of your experiment. Instructions for creating this file are available as part of the profiling-recipe^42^ and profiling-handbook.
48. Follow the instructions in the profiling-template^43^ and profiling-handbook to create a new template analysis repository and weld a copy of the profiling-recipe into it.
49. Follow the instructions in the profiling-recipe to set up your profiling configuration file for your current batch of data.
50. Execute the profiling-recipe script to create annotated, normalized, and/or feature reduced profiles. This step will take a few minutes per plate.

### Data analysis

**TIMING** variable

51. Use the per-well profiles to analyze patterns in the data. How to do so is an area of active research and is customized to the biological question at hand. A typical profiling data analysis workflow begins with the per-well profiles; for most applications, a key step is measuring the similarity (or, equivalently, distance) between each sample’s profile and all other profiles in the experiment. Methods often used for measuring similarity or distance are Pearson’s correlation, Spearman’s correlation, Euclidean distance, and cosine distance. For QC purposes, it is customary to check that replicates of the same sample yield small distances. If positive controls are available (that is, samples that are known to yield similar phenotypes), their replicates can also be checked for producing small distances relative to random pairs of samples. Samples are often clustered using hierarchical clustering, although other clustering methods may also be used. A discussion of best practices in analyzing phenotypic profiling data (including Cell Painting) can be found in Caicedo et al 2017^44^.

## Troubleshooting

Troubleshooting advice can be found in Table 4. We also recommend the Assay Guidance Manual^45^ as an excellent source for learning about and troubleshooting high content imaging assays of this type.

## Timing

**Steps 1–9, cell culture:** it typically takes ∼2–3 d for the cells to reach appropriate confluency, depending on cell type and growth conditions. Harvesting the cells (Steps 3–7) takes 20 min, and seeding the cells (Step 8) takes 20 min. After seeding, the cells should be cultured for 2–5 d before staining.

**Step 10(A), addition of a small-molecule library (optional):** variable; approximately 2–3 d for one batch experiment of 384-well plates. Addition of a compound library takes ∼1 h for one batch experiment of 384-well plates, including reagent preparation and media change, and 1–2 d for compound incubation.

**Step 10(B), CRISPR knockdown (optional):** variable; approximately 4–6 d for one batch experiment of 384-well plates. Addition of polybrene and lentiviral CRISPR library takes ∼2 h for one batch experiment of 384-well plates, including centrifugation. Media change(s) take ∼1 h for one batch experiment of 384-well plates, and an additional 3-5 d for incubation.

**Step 10(C), ORF overexpression (optional):** variable; approximately 2–4 d for one batch experiment of 384-well plates. Addition of polybrene and lentiviral ORF library takes ∼2 h for one batch experiment of 384-well plates, including centrifugation. Media change takes ∼1 h for one batch experiment of 384-well plates, and an additional 1-2 d for incubation.

**Steps 11–21, staining and fixation:** ∼2.5–3 h including reagent preparation for one batch experiment of 384-well plates. The total timing will vary depending on the number of plates in the experiment and the automation available. We have found that up to 12 plates can be simultaneously fixed and stained as one batch in this span of time. We recommend having all staining and fixation solutions be prepared before beginning mitotracker staining to allow for sufficient time to prepare equipment during incubation steps.

**Steps 22–24, automated image acquisition:** variable; ∼1.5 h per 384-well plate, for nine fields of view per well and typical exposure times (and as little as 1 h per plate for smaller numbers of fields of view). The total time varies depending on the number of sites imaged per plate, the number of channels acquired per site, the number of Z planes each channel is imaged in, and the exposure time for each channel.

**Steps 25–44, morphological image feature extraction from microscopy data):** variable, very dependent on computing setup; 24 h - 1 week per 384-well plate. It takes ∼10 min per plate for CellProfiler to scan the images in the input folder(s) after manually dragging/dropping them into the CellProfiler interface. Optimizing the segmentation pipeline before running it can take from minutes to hours depending on the diversity of the input images and how different they are from the starting conditions.

The pipeline execution time will depend on the computing setup; run times on a single computing node of 20 s (illumination correction), 20 s (segmentation), 30 s (QC), and 10 min (analysis) per field of view are typical; these equate to about 650 CPU-hours of processing per plate, most which can be fully parallelized. Handling of QC results, once a visualization tool like the open-source KNIME is set up, takes 20 s per plate (see Box 2). A substantial time saving can be achieved if you run the feature extraction, segmentation, and QC pipelines on a distributed computing cluster, which massively parallelizes the processing as compared with running it on a single local computer, as well as removing slowdowns in CellProfiler that can accumulate after several hundreds of images are processed in a single run.

**Steps 45–50, Creation, normalization, and feature reduction of per-well profiles:** variable; collation of CSVs (if CellProfiler is run in parallel with ExportToSpreadsheet rather than on a single machine with ExportToDatabase) and aggregation into per-well mean profiles can take 8-16 unattended hours per 384-well plate; plates may be run in parallel. Metadata CSV creation and repository setup will take ∼30 minutes for an experienced user and may take 2-3 hours for an inexperienced user. Execution of the profiling recipe steps takes <5 min of processing time per 384-well plate.

**Step 51, data analysis:** variable; ∼1 h for basic analysis of replicate quality and signature strength. Time for additional analysis varies substantially depending on the problem at hand.

## Anticipated Results

The automated imaging protocol will produce a large number of acquired single-channel images in 16-bit TIF format; each resultant image will be ∼1,000 × 1,000 pixels and ∼2.5 MB in size. The total number of images generated equals (number of samples tested) × (number of sites imaged per well) × (channels imaged). In terms of data storage, a single 384-well microplate will produce 3,456 fields of view, or 17,280 images total across all channels, for a total of ∼40 GB per microplate. Each additional brightfield channel acquired adds another 3,456 images to the total image count and thus an additional ∼8GB of storage.

The illumination correction pipeline will yield an illumination correction image per plate and channel. Each image is ∼4.5 MB so one microplate’s worth of illumination correction images with 5 fluorescent channels and 1 brightfield channel will occupy ∼26.5 MB of storage space.

The optional QC pipeline will produce a set of numerical measurements extracted at the image level, and will export them to a series of .csv’s. These measurements can optionally be used for comparing quality between batches or for flagging images that you may want to remove from downstream processing because of focal blur or saturation artifacts.

The segmentation pipeline outputs a single image for each well with segmented nuclei and cell objects overlaid onto the images. The quality of the image features extracted by the analysis pipeline and downstream profiling will depend on accurate nuclei and cell body segmentation determined in the segmentation pipeline. First, the nuclei are identified from the Hoechst image because it is a high-contrast stain for a well-separated organelle; subsequently, the nucleus, along with an appropriate channel, is used to delineate the cell body. We have found that the SYTO 14 image is the most amenable for finding cell edges, as it has fairly distinct boundaries between touching cells. For help in optimizing the output, if needed, see the TROUBLESHOOTING table (Table 4).

The image analysis pipeline produces the raw numerical features extracted from nuclei, cell, and cytoplasm objects identified in the images. If run locally, these are saved as SQLite database files with one table per object. If run distributed, these are saved into per-object .csvs (i.e. Cells.csv, Cytoplasm.csv, and Nuclei.csv) in a per-site folder. The .csv’s contain one row for each object in each image, and ∼2,000 columns containing the values for the different morphological features that have been measured for that object. The pipeline also outputs Experiment.csv and Image.csv containing information about the experiment and whole image measurements, respectively. The pipeline also saves cell and nuclei object outlines, though these can optionally be toggled off. Generally, the pipeline is not configured to save any processed images (to conserve data storage space), but additional SaveImages modules can be used for this purpose if desired. The combination of data tables and object outlines from the feature extraction pipeline is typically ∼5 MB per site.

After running the profiling scripts to create the per-well profiles, the output will be a number of image-based profile files in CSV format. Each row of this file represents a data vector for an individual plate and well, with each column containing a per-well mean measurement of a given image feature. Initial profiles will contain 6,000 columns of raw features; normalized profile files will contain the same number of columns but will have had each column independently normalized to the median and MAD of the data distribution.

## Data availability

All images, single cell profiles, and processed profiles are available at the Cell Painting Gallery at https://registry.opendata.aws/cellpainting-gallery/ under accession numbers cpg0000-jump-pilot and cpg0001-cellpainting-protocol. Processed profiles, metadata, and a detailed description of all plates are available in the paper’s GitHub repository (https://github.com/carpenterlab/2022_Cimini_NatureProtocols) and the data repositories found therein (https://github.com/jump-cellpainting/pilot-cpjump1-data, https://github.com/jump-cellpainting/pilot-cpjump1-fov-data, and https://github.com/jump-cellpainting/pilot-data-public).

## Code availability

All code necessary to reproduce these analyses is available in the paper’s GitHub repository (https://github.com/carpenterlab/2022_Cimini_NatureProtocols) and additionally archived to Zenodo (https://doi.org/10.5281/zenodo.7267354).

## Acknowledgments

The authors appreciate the more than 100 scientists who have contributed to the organization and scientific direction of the JUMP Cell Painting Consortium. The authors thank the original developers of earlier versions of the protocol, who contributed to prior papers describing the assay ^3,4^; those who are not also authors on this paper are: M.A Bray, H. Han, C.T. Davis, B. Borgeson, C. Hartland, S.M. Gustafsdottir, C.C. Gibson, V. Ljosa, K.L. Sokolnicki, J.A. Wilson, D. Walpita, M.M. Kemp, K. Petri Seiler, H.A. Carrel, T.R. Golub, S.L. Schreiber, P.A. Clemons, and A.F. Shamji.

## Author Contributions

BAC, SNC, MK-A, LM, AG, BF, SG, JBC C-HL, EM, SS, JDB, TG, DG, TM, JP, AS,SES, AV, GW, SY, BZ, and AEC contributed to experimental design. MK-A, LM, AG, BF, PB, JBC, EM, PJA, JDB, TG, DH, GH, KJ, MM, LM, TM, JP, AS, AV, GW, and SY conducted laboratory experiments. BAC, SNC, DJL, NJ, HSA, SH, LM, GW, and BZ performed image and/or data analysis. BAC, SNC, MK-A, LM, PB, SG, JBC, C-HL, EM, SS, JDB, GH, MM, LM, TM, AS, SES, GW, BZ, and AEC contributed to data interpretation. BAC, DJL, PJA, SH, EW, and AEC contributed to protocol development. BAC, SNC, MK-A, LM, AG, BF, SG, JBC, SS, PJA, DG, EW, and AEC wrote the manuscript, with all authors contributing to editing and giving manuscript approval. Supervision and administration were carried out by BAC, SS, and AEC.

## Boxes

### Box 1

Configuration of the pipelines for batch processing on a computer cluster

We recommend using a computing cluster, such as a high-performance server farm, or a cloud-computing platform such as Amazon Web Services (AWS) for analyzing Cell Painting experiments to speed processing, especially for experiments with >1,000 fields of view. The typical batch processing workflow is to distribute smaller subsets of the acquired images to run on individual computing nodes. Each subset is run using CellProfiler in ‘headless’ mode—i.e., from the command line without the user interface. The headless runs are executed in parallel, with a concomitant decrease in overall processing time. Initial preparation to run on a cluster or cloud requires expert setup and will probably require enlisting the help of your IT department or local cloud computing expert. Please refer to our Distributed CellProfiler GitHub repository wiki (https://github.com/CellProfiler/Distributed-CellProfiler/wiki) for more details on cloud computing using AWS and our command line startup guide https://github.com/CellProfiler/CellProfiler/wiki/Getting-started-using-CellProfiler-from-the-command-line for more details on cluster computing. Additional information on batch processing on a cluster or in the cloud is available on our video Headless CellProfiler/DistributedCellProfiler Tutorial (https://youtu.be/LuJxIGGhRek).

When running batch processes, we recommend exporting data to spreadsheets as it is easier to aggregate a large number of spreadsheets than databases. Uncheck or remove any ExportToDatabase modules in your pipelines and add instead ExportToSpreadsheet modules.

There are two ways to create batch information for CellProfiler: LoadData and CreateBatchFiles.

**LoadData:** LoadData is our preferred method, especially for use with Distributed CellProfiler and cloud computing. Detailed instructions can be found at https://cytomining.github.io/profiling-handbook/

1. Insert the LoadData module into the pipeline by pressing the ‘+’ button and selecting the module from the ‘File Processing’ category. Move this module to the beginning of the pipeline and configure it to find your file.
2. Create LoadData.csvs. If you are using a Perkin Elmer microscope you can use our pe2loaddata script, available at https://github.com/broadinstitute/pe2loaddata, to create the LoadData.csv file. Alternatively, if you have loaded your images into CellProfiler using the Input modules, you can export a file list from CellProfiler using File > Export > Image Set Listing.
3. The end result of this step will be a LoadData.csv file. This file, when used in conjunction with a CellProfiler pipeline (.cppipe), contains the information needed to run in ‘headless’ mode on the cluster or in the cloud

**CreateBatchFiles:** If you have already created a CellProfiler project (.cpproj) with fully populated Input modules, you can

1. Insert the CreateBatchFiles module into the pipeline by pressing the ‘+’ button and selecting the module from the ‘File Processing’ category. Move this module to the end of the pipeline.
2. Configure the CreateBatchFiles module by setting the ‘Local root path’ and ‘Cluster root path’ settings. If your computer mounts the file system differently than the cluster computers, CreateBatchFiles can replace the necessary parts of the paths to the image and output files. For instance, a Windows machine might access files images by mounting the file system using a drive letter, e.g., `C:\your_data\images` and the cluster computers access the same file system using `/server_name/your_name/your_data/images`. In this case, the local root path is `C:\` and the cluster root path is `/server_name/your_name.` You can press the ‘Check paths’ button to confirm that the path mapping is correct.
3. Press the ‘Analyze Images’ button at the bottom-left of the interface.
4. The end result of this step will be a ‘Batch_data.h5’ (HDF5 format) file. This file contains the pipeline plus all information needed to run on the cluster.
5. This file will be used as input to CellProfiler on the command line, in order for CellProfiler to run in ‘headless’ mode on the cluster or in the cloud. There are a number of command-line arguments to CellProfiler that allow customization of the input and output folder locations, as well as which images are to be processed on a given computing node. Enlist an IT specialist to specify the mechanism for sending out the individual CellProfiler processes to the computing cluster nodes. Please refer to our GitHub webpage https://github.com/CellProfiler/CellProfiler/wiki/Adapting-CellProfiler-to-a-LIMS-environment for more details.

### Box 2

Quantifying Quality Control with CellProfiler

Though quantifying the quality of your images is not strictly necessary to obtain profiles, you may find it critical to ensuring your data is of high enough staining and imaging quality to measure your phenotypes of interest; running it alongside each batch of data can help discover workflow issues that can be addressed before subsequent runs. You can integrate these quality measurements into your analysis pipeline or you can run them as a separate pipeline before analysis, as we have in this protocol.

1. Start CellProfiler.
2. Load the qc pipeline into CellProfiler by selecting File > Import > Pipeline from File from the CellProfiler main menu and then selecting qc.cppipe or dragging and dropping the pipeline from your file browser into the pipeline panel on the left of the interface.
3. Select the Images input module in the ‘Input modules’ panel to the top-left of the interface. From your file browser, drag and drop the folder(s) containing your raw images into the ‘File list’ panel. See the Computational Equipment section for a link to raw image files that can be used as an example in this protocol.
4. Click the ‘Output settings’ button at the bottom-left of the interface. Select an appropriate ‘Default Output Folder’ into which the qc measurements will be saved.
5. Save the current settings to a project (.cpproj) file containing the pipeline, the list of input images, and the output location by selecting File > Save Project. Enter the desired project filename in the dialog box that appears. This is not necessary for the running of this step, however it functions as a snapshot of your complete setup, allowing you to directly replicate your work by loading this .cpproj instead of the .cppipe you started with.
6. Press the ‘Analyze Images’ button at the bottom-left of the interface. A progress bar in the bottom-right will indicate the estimated time of completion.
7. Visualize the output from the CellProfiler QC pipeline; our recommendation is to use KNIME, an open-source data analytics tool.
  a. Download and install the current version of KNIME (https://www.knime.com/downloads)
  b. Within the KNIME client, install the HCS Tools extension (https://www.knime.com/community/hcs-tools)
  c. Download the KNIME workflow JUMP_QC_Plate-CV_v1.knwf from https://github.com/broadinstitute/imaging-platform-pipelines/tree/master/JUMP_production
8. Run KNIME and load KNIME workflow
  a. Right-click the CSV Reader Node “Top Line Per-Image”, select “Configure…”, and set the File path to the TopLineImage.csv output from the CellProfiler workflow
  b. Right-click the CSV Reader Node “Top Line Per-Object”, select “Configure…”, and set the File path to the TopLineCells.csv output from the CellProfiler workflow
  c. Run the KNIME workflow (Green, double-arrow button in the menu bar)
  d. Inspect the plots, and right-click the final nodes (right-click, select “View: …”

## References

1. Chandrasekaran, S. N., Ceulemans, H., Boyd, J. D. & Carpenter, A. E. Image-based profiling for drug discovery: due for a machine-learning upgrade? Nat. Rev. Drug Discov. 20, 145–159 (2021).

2. Pratapa, A., Doron, M. & Caicedo, J. C. Image-based cell phenotyping with deep learning. Curr. Opin. Chem. Biol. 65, 9–17 (2021).

3. Bray, M.-A. et al. Cell Painting, a high-content image-based assay for morphological profiling using multiplexed fluorescent dyes. Nat. Protoc. 11, 1757–1774 (2016).

4. Gustafsdottir, S. M. et al. Multiplex cytological profiling assay to measure diverse cellular states. PLoS One 8, e80999 (2013).

5. Caicedo, J. C. et al. Cell Painting predicts impact of lung cancer variants. Mol. Biol. Cell mbcE21110538 (2022).

6. Heiser, K. et al. Identification of potential treatments for COVID-19 through artificial intelligence-enabled phenomic analysis of human cells infected with SARS-CoV-2. bioRxiv (2020) doi:10.1101/2020.04.21.054387.

7. Nyffeler, J. et al. Bioactivity screening of environmental chemicals using imaging-based high-throughput phenotypic profiling. Toxicol. Appl. Pharmacol. 389, 114876 (2020).

8. Carey, K. L. et al. TFEB Transcriptional Responses Reveal Negative Feedback by BHLHE40 and BHLHE41. Cell Rep. 33, 108371 (2020).

9. Strobel, S. et al. Discovering cellular programs of intrinsic and extrinsic drivers of metabolic traits using LipocyteProfiler. bioRxiv (2021).

10. Simm, J. et al. Repurposing High-Throughput Image Assays Enables Biological Activity Prediction for Drug Discovery. Cell Chem Biol 25, 611–618.e3 (2018).

11. Moshkov, N. et al. Predicting compound activity from phenotypic profiles and chemical structures. bioRxiv (2022) doi:10.1101/2020.12.15.422887.

12. Rohban, M. H. et al. Virtual screening for small-molecule pathway regulators by image-profile matching. Cell Syst 13, 724–736.e9 (2022).

13. Chandrasekaran, S. N. et al. Three million images and morphological profiles of cells treated with matched chemical and genetic perturbations. bioRxiv 2022.01.05.475090 (2022) doi:10.1101/2022.01.05.475090.

14. Way, G. P. et al. Morphology and gene expression profiling provide complementary information for mapping cell state. Cell Systems 0, (2022).

15. Haghighi, M., Singh, S., Caicedo, J. & Carpenter, A. High-Dimensional Gene Expression and Morphology Profiles of Cells across 28,000 Genetic and Chemical Perturbations. bioRxiv 2021.09.08.459417 (2021) doi:10.1101/2021.09.08.459417.

16. Caicedo, J. C. et al. Nucleus segmentation across imaging experiments: the 2018 Data Science Bowl. Nat. Methods 16, 1247–1253 (2019).

17. Dobson, E. T. A. et al. ImageJ and CellProfiler: Complements in Open-Source Bioimage Analysis. Current Protocols 1, e89 (2021).

18. Schmidt, U., Weigert, M., Broaddus, C. & Myers, G. Cell Detection with Star-Convex Polygons. in Medical Image Computing and Computer Assisted Intervention –MICCAI 2018 265–273 (Springer International Publishing, 2018).

19. Stringer, C., Wang, T., Michaelos, M. & Pachitariu, M. Cellpose: a generalist algorithm for cellular segmentation. Nat. Methods 18, 100–106 (2021).

20. Stirling, D. R. et al. CellProfiler 4: improvements in speed, utility and usability. BMC Bioinformatics 22, 1–11 (2021).

21. Rohban, M. H. et al. Systematic morphological profiling of human gene and allele function via Cell Painting. Elife 6, (2017).

22. Cross-Zamirski, J. O. et al. Label-free prediction of cell painting from brightfield images. Sci. Rep. 12, 10001 (2022).

23. Van Rossum, G. & Drake, F. L. Python 3 Reference Manual: (Python Documentation Manual Part 2). (CreateSpace Independent Publishing Platform, 2009).

24. Berthold, M. R. et al. KNIME: The Konstanz Information Miner. in Studies in Classification, Data Analysis, and Knowledge Organization (GfKL 2007) (Springer, 2007).

25. Stöter, M. et al./person-group>. CellProfiler and KNIME: Open Source Tools for High Content Screening. in Target Identification and Validation in Drug Discovery: Methods and Protocols (eds. Moll, J. & Colombo, R.) 105–122 (Humana Press, 2013).

26. Singh, S., Bray, M.-A., Jones, T. R. & Carpenter, A. E. Pipeline for illumination correction of images for high-throughput microscopy. J. Microsc. 256, 231–236 (2014).

27. Schindelin, J. et al. Fiji: an open-source platform for biological-image analysis. Nat. Methods 9, 676–682 (2012).

28. Schindelin, J., Rueden, C. T., Hiner, M. C. & Eliceiri, K. W. The ImageJ ecosystem: An open platform for biomedical image analysis. Mol. Reprod. Dev. 82, 518–529 (2015).

29. Brocher, J. biovoxxel/BioVoxxel-Figure-Tools: BioVoxxel-Figure-Tools_1.2.1b. (2022). doi:10.5281/zenodo.7268128.

30. Cimini, B. A. et al. Broad Institute Imaging Platform Profiling Handbook. https://github.com/cytomining/profiling-handbook.

31. Reback, J. et al. pandas-dev/pandas: Pandas 1.3.4. (2021). doi:10.5281/zenodo.5574486.

32. Harris, C. R. et al. Array programming with NumPy. Nature 585, 357–362 (2020).

33. Hunter, J. D. Matplotlib: A 2D Graphics Environment. Computing in Science Engineering 9, 90–95 (2007).

34. Waskom, M. seaborn: statistical data visualization. J. Open Source Softw. 6, 3021 (2021).

35. van der Walt, S. et al. scikit-image: image processing in Python. PeerJ 2, e453 (2014).

36. Pedregosa, F. et al. Scikit-learn: Machine Learning in Python. J. Mach. Learn. Res. 12, 2825–2830 (2011).

37. Satopaa, V., Albrecht, J., Irwin, D. & Raghavan, B. Finding a ‘Kneedle’ in a Haystack: Detecting Knee Points in System Behavior. in 2011 31st International Conference on Distributed Computing Systems Workshops 166–171 (2011).

38. Kluyver, T. et al./person-group>. Jupyter Notebooks -- a publishing format for reproducible computational workflows. in Positioning and Power in Academic Publishing: Players, Agents and Agendas (eds. Loizides, F. & Schmidt, B.) 87–90 (IOS Press, 2016).

39. Way, G. et al. Pycytominer: Data processing functions for profiling perturbations.

40. Singh, S. et al. cytominer-database.

41. Tange, O. GNU Parallel 2018. (Lulu.com, 2018).

42. Chandrasekaran, S. N., Weisbart, E., Way, G., Carpenter, A. & Singh, S. Broad Institute Imaging Platform Profiling Recipe.

43. Chandrasekaran, S. N., Way, G., Carpenter, A. & Singh, S. Broad Institute Imaging Platform Profiling Template.

44. Caicedo, J. C. et al. Data-analysis strategies for image-based cell profiling. Nat. Methods 14, 849–863 (2017).

45. Assay Guidance Manual. (Eli Lilly & Company and the National Center for AdvancingTranslational Sciences, 2012).

