## Extended Data and Supplemental Figures for "Optimizing the Cell Painting assay for image-based profiling"

### Extended Data Figures

JUMP-MOA Plate - Compound Number

| | 1 | 2 | 3 | 4 | 5 | 6 | 7 | 8 | 9 | 10 | 11 | 12 | 13 | 14 | 15 | 16 | 17 | 18 | 19 | 20 | 21 | 22 | 23 | 24 |
|---|---|---|---|---|---|---|---|---|---|---|---|---|---|---|---|---|---|---|---|---|---|---|---|---|
| A | 63 | 69 | 27 | 51 | 33 | 52 | 19 | 6 | 20 | 76 | | 48 | 77 | 15 | 49 | 81 | 84 | 74 | 55 | 36 | 90 | 75 | 67 | 35 |
| B | 16 | 32 | 81 | 47 | 31 | 15 | 45 | 57 | 30 | 21 | 53 | 31 | 42 | 65 | 52 | 44 | 59 | | 23 | 66 | 56 | 28 | 50 | 64 |
| C | 58 | 66 | 43 | 9 | 72 | 83 | 38 | 4 | 65 | 62 | 12 | 82 | 61 | 37 | 89 | 22 | 11 | 60 | 73 | 47 | 53 | 25 | 39 | 71 |
| D | 15 | 30 | 87 | 39 | 74 | 71 | 36 | 5 | 68 | 7 | 35 | 73 | 64 | 54 | 24 | 38 | | 33 | | 83 | 83 | 42 | 58 | 86 |
| E | 64 | 64 | 70 | 70 | 13 | 13 | | | 54 | 54 | 50 | 50 | 5 | 77 | 7 | 49 | 41 | 84 | 68 | 55 | 79 | 90 | 80 | 67 |
| F | 57 | 68 | 11 | 61 | 32 | 25 | | 81 | 90 | 49 | 39 | 87 | 43 | 13 | 62 | 23 | 66 | 20 | 2 | 34 | 51 | 62 | 12 | |
| G | 66 | 58 | 9 | 43 | 83 | 72 | 4 | 38 | 62 | 65 | 82 | 12 | 3 | 46 | 14 | 57 | 29 | 16 | 32 | | 31 | 85 | | 30 |
| H | 67 | 89 | 41 | 35 | 55 | 40 | 89 | 26 | 21 | | 22 | 46 | 88 | 51 | 34 | 78 | 63 | 4 | 65 | 58 | 38 | 1 | 76 | 52 |
| I | 69 | 63 | 51 | 27 | 52 | 33 | 6 | 19 | 76 | 20 | 48 | | | 61 | 40 | 89 | 21 | 11 | 45 | 73 | 26 | 53 | 87 | 39 |
| J | | 90 | 46 | 22 | 5 | 45 | 47 | 36 | 29 | 67 | 79 | 37 | 86 | 8 | 70 | 56 | 69 | 6 | 19 | 10 | | 24 | 78 | 18 |
| K | 88 | 2 | 18 | 1 | 17 | 44 | 23 | 56 | 86 | 28 | 42 | 10 | 15 | 3 | 81 | 14 | 74 | 29 | 36 | 32 | 75 | 31 | 35 | |
| L | 37 | 29 | 7 | 3 | 60 | 79 | 3 | 60 | 25 | 84 | 71 | 41 | 18 | 72 | 10 | 76 | 9 | 70 | 54 | 63 | 72 | 27 | 4 | 43 |
| M | 24 | 24 | 34 | 34 | | | 59 | 59 | 8 | 8 | 78 | 78 | 37 | 5 | 22 | 7 | 60 | 41 | 47 | 68 | 25 | 79 | 71 | 80 |
| N | 80 | 53 | 73 | 74 | 26 | | 85 | 11 | 61 | 75 | 40 | 80 | 48 | 12 | 17 | 50 | 1 | 17 | 13 | 2 | 6 | | 33 | 59 |
| O | 2 | 88 | 1 | 18 | 44 | 17 | 56 | 23 | 28 | 86 | 10 | 42 | 46 | | 57 | 40 | 16 | 21 | | 45 | 85 | 26 | 30 | 87 |
| P | 77 | | 49 | 85 | 84 | 55 | 14 | 14 | 75 | 16 | | 77 | 27 | 82 | 82 | 48 | 28 | 9 | 44 | 19 | 8 | 69 | 20 | 88 |

Extended Data Figure 1 - The JUMP-MOA compound plate map: unlabeled wells are DMSO only, all other wells are labeled to show distribution of compound replicates across the entire plate. A version of this figure grouped by MOA rather than compound is available as Figure 2A.

a)

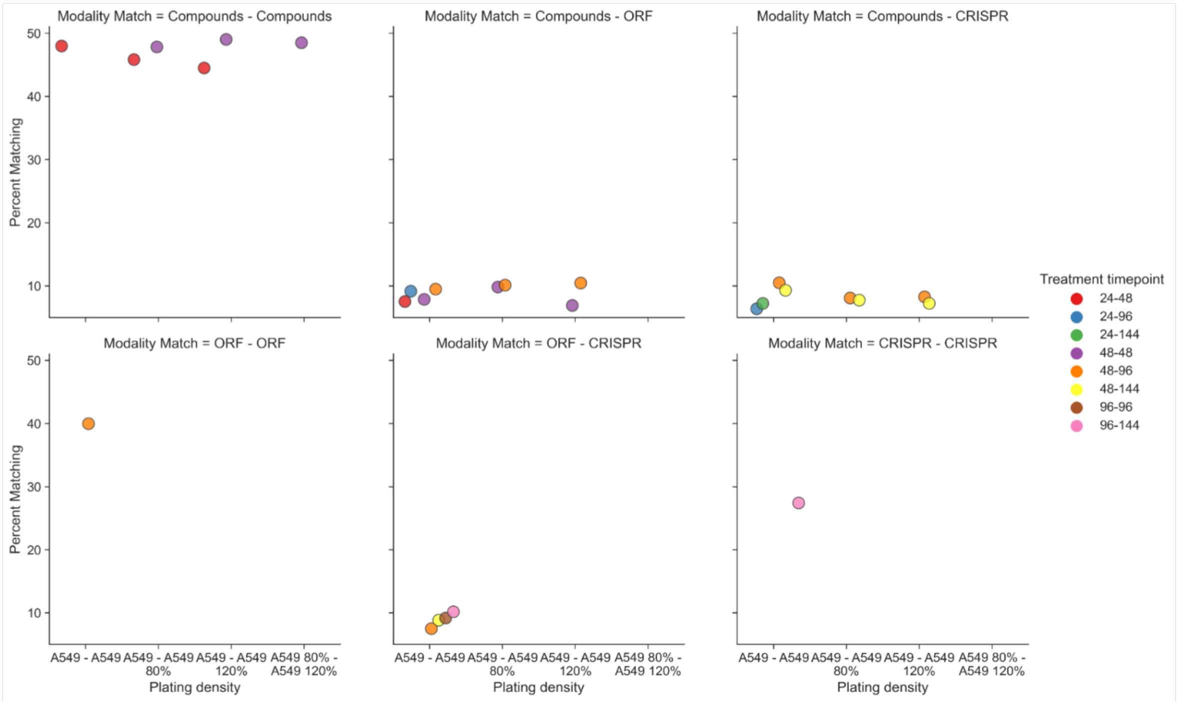


b)

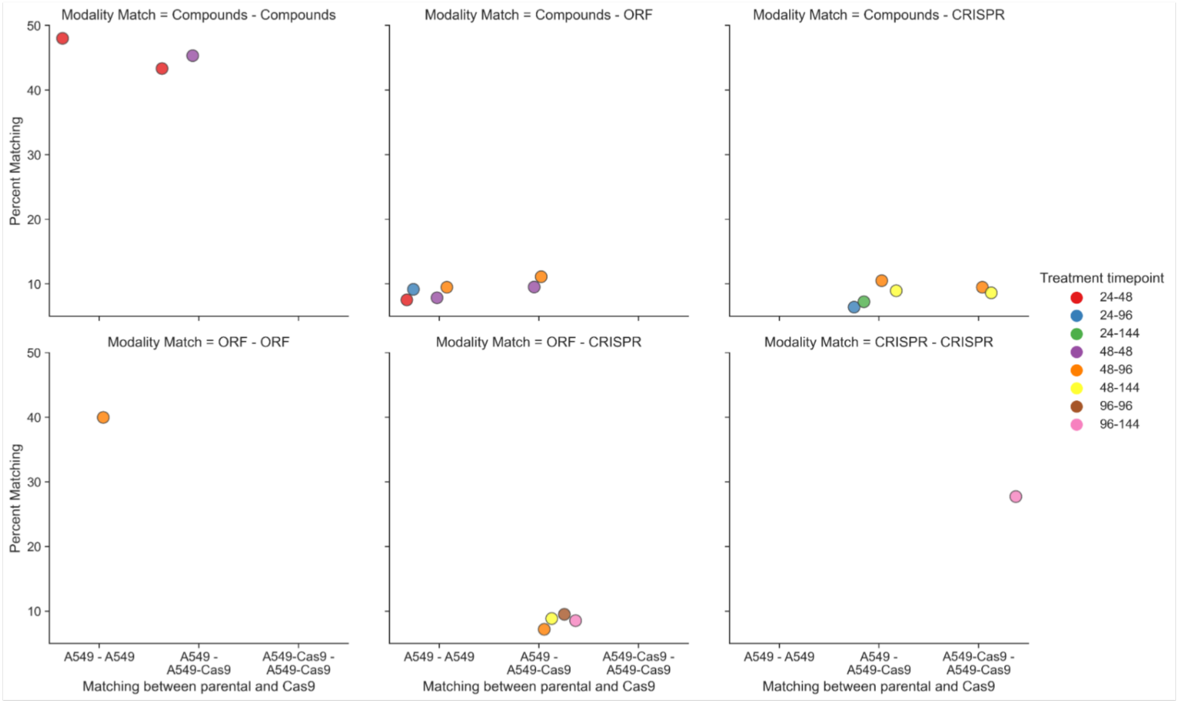


Extended Data Figure 2- A) Assessment of decreasing or increasing the number of cells plated by 20% on ability to perform Percent Matching. Few consistent effects are seen. B) Assessment of using parental cell lines or Cas9 expressing polyclonal lines for compound treatments. Few consistent effects are seen; ability to match compounds in parental vs Cas9 cells against CRISPR (required to be performed in Cas9 cells) does not seem extremely different at either common time point.

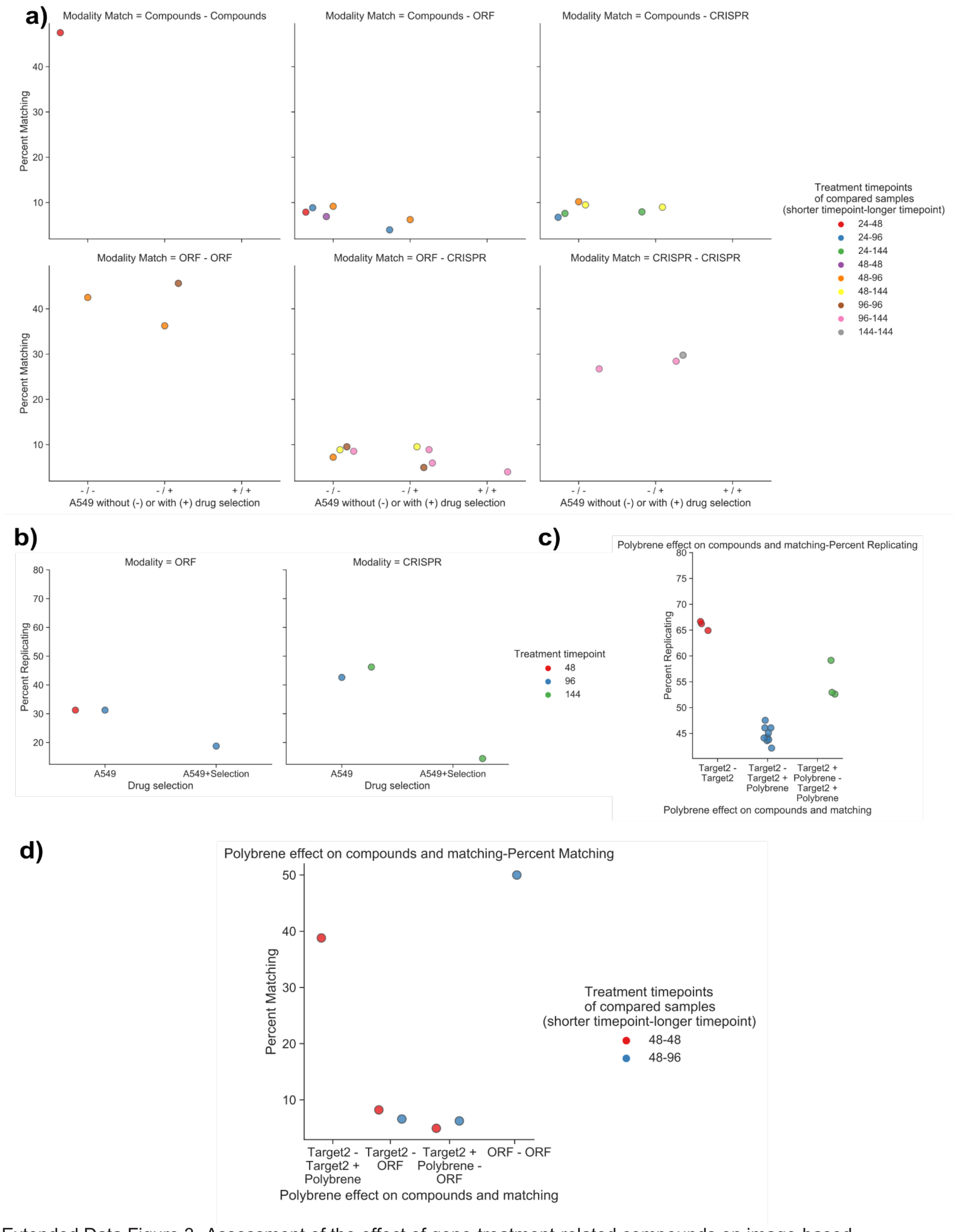


Extended Data Figure 3- Assessment of the effect of gene-treatment-related compounds on image-based profiling. A) Addition of blasticidin to ORF plates or puromycin to CRISPR plates does not appear to improve percent matching across modalities vs unselected plates. B) Addition of selection compounds may have a deleterious effect on percent matching vs unselected plates, though we cannot rule out that this is due to fewer replicates for the selected plates than the unselected ones. C) Addition of 4µg/mL polybrene for 24 hours may produce a phenotypic effect; polybrene addition displays decreased inter-treatment cross-plate percent replicating vs intra-treatment cross-plate-percent replicating, even though both sets of plates were part of the same batch. D) Addition of polybrene to Target2 treated cells does not improve percent matching between Target2 treated plates and ORF treated plates from a previous batch.

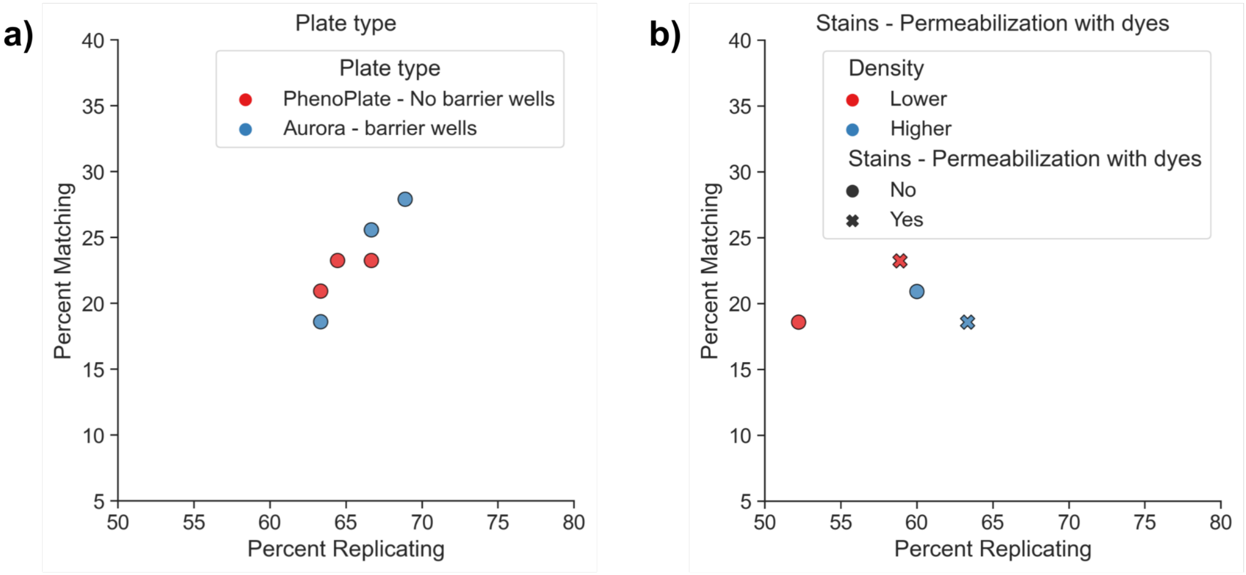


Extended Data Figure 4- A) PhenoPlates without barrier wells and Aurora plates with barrier wells produce comparable percent replicating and percent matching results. B) Performing permeabilization at the same time as staining produces comparable percent replicating and percent matching results.

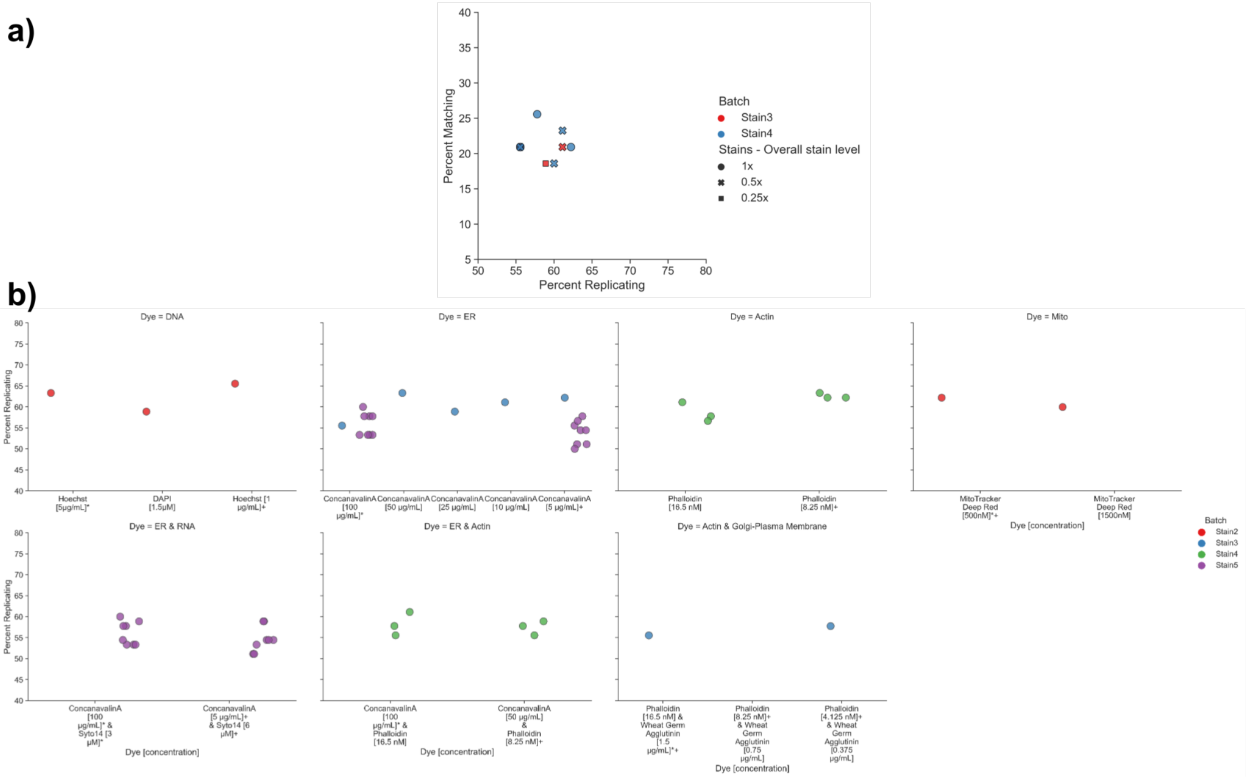

Extended Data Figure 5- A) Within batches, reducing all dyes by 2 or 4 fold produces comparable percent replicating and comparable but perhaps slightly decreased percent matching results. B) All quantitatively tested stain concentration changes, broken out by the dye(s) perturbed.

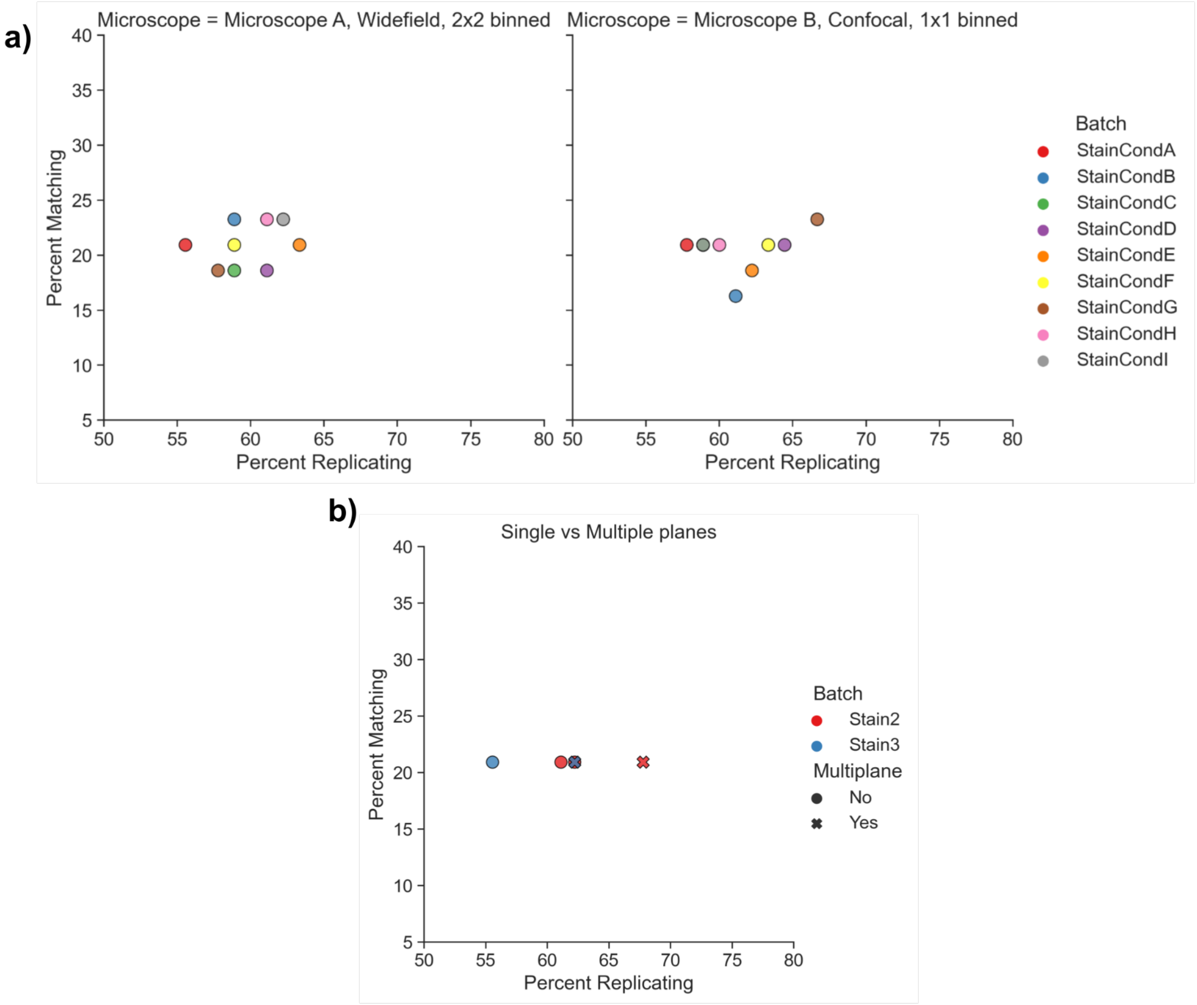


Extended Data Figure 6- A) Imaging of the same plates on a widefield microscope with 2x2 binning vs a different manufacturer's microscope in confocal with 1x1 binning. No major differences are observed. B) Acquisition of multiple Z planes slightly improves percent replicating but not percent matching in two batches.

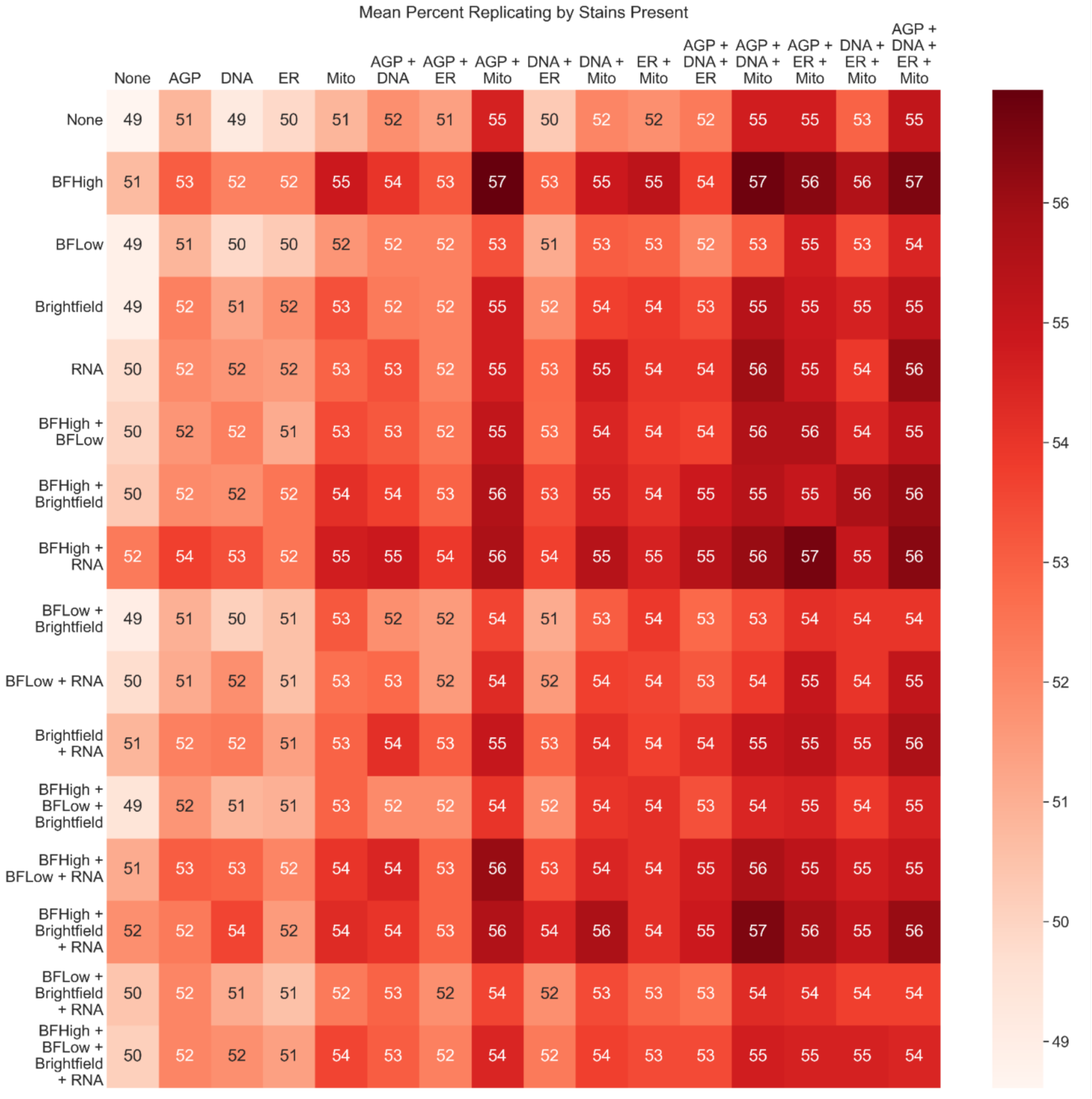


Extended Data Figure 7 - Mean percent replicating of eight JUMP-MOA plates stained with the final staining conditions after dropping out all feature names containing individual channel names before performing feature selection and calculation of percent replicating. “None” means that only AreaShape features are present. To create a sufficiently compact data representation, theeightchannels present were split 4 each onto the X (AGP, DNA, ER, and Mito) and Y (RNA, Brightfield, BFHigh, and BFLow) axes; this allows visualization of the 256 possible unique combinations. Note that these results are not the same as truly having the stains not present, as a) a channel still may have been used in creating the initial segmentation results and b) it does not account for cross talk between stains. An alternate representation of this data is presented in Extended Data Figure 8.

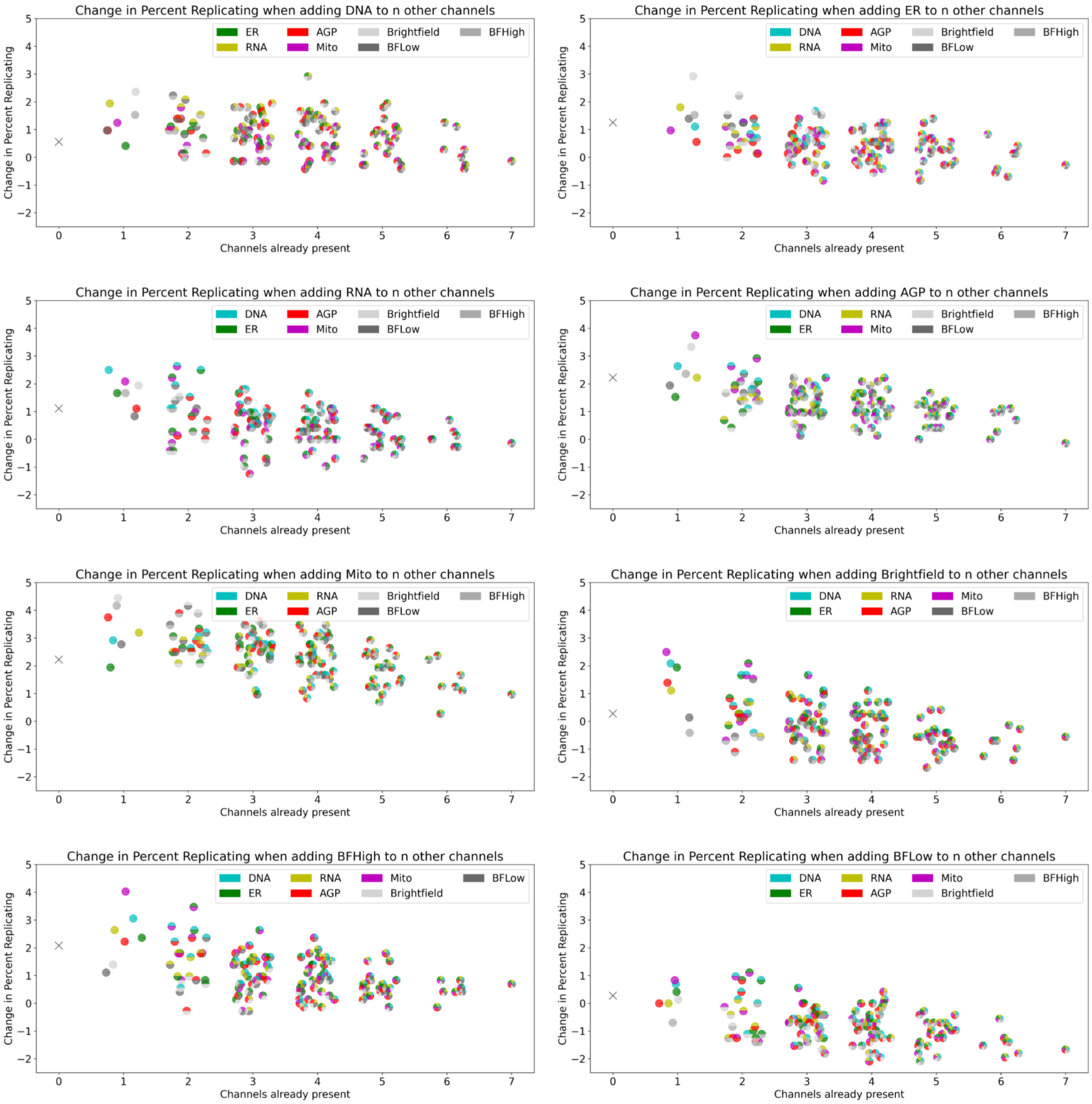


Extended Data Figure 8- Change in mean percent replicating of eightJUMP-MOA plates stained with the final staining conditions when an individual stain is present vs not; each chart shows for a particular stain when added to the non-channel-specific features plus zero or more other stains (x axis) the change in the mean percent replicating (y axis) when those features are included. The color(s) of each marker indicate which channel(s) is/are already present. Note that these results are not the same as truly having the stains not present, as a) a channel still may have been used in creating the initial segmentation results and b) it does not account for cross talk between stains. The absolute percent replicating numbers are available in Extended Data Figure 7.

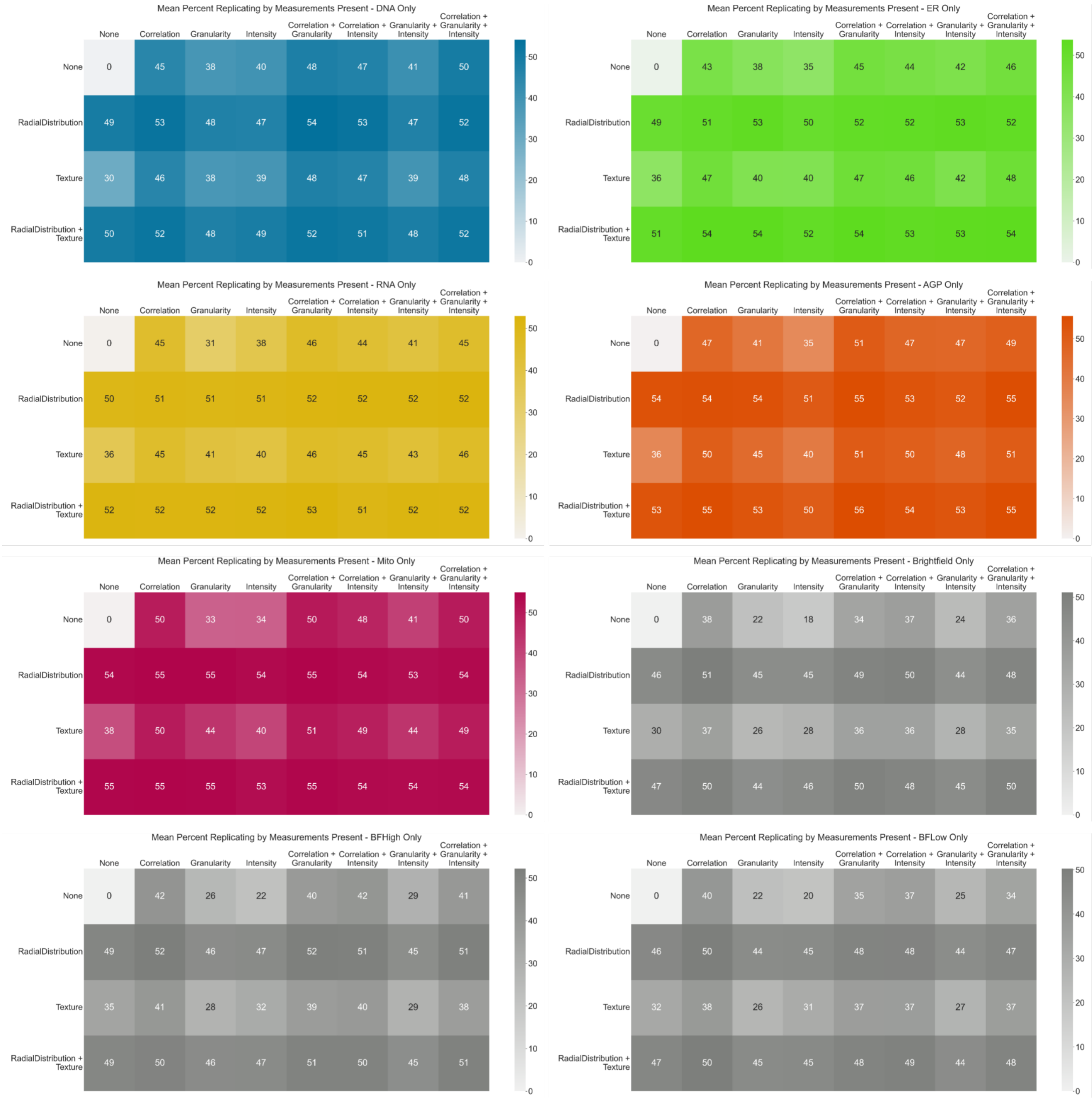


Extended Data Figure 9 - Stain-by-stain breakdown of mean percent replicating of eightJUMP-MOA plates stained with the final staining conditions after dropping out all possible combinations of features from the 5 stain-specific feature categories before performing feature selection and calculation of percent replicating. Unlike the analyses in Extended Data Figure 7 and Extended Data Figure 8, non-channel-specific feature categories (AreaShape and Neighbors) are not included here, as the goal is to assess the information contribution of each feature category in each stain. To create a sufficiently compact data representation, the 5 categories present were split 3 each onto the X (Correlation, Granularity, and Intensity) and2 onto the Y (RadialDistribution, and Texture) axes; this allows visualization of the 31 possible unique combinations.

### Supplementary Information

| Cell Line | Clone Type | CRISPR Efficiency |
|---|---|---|
| U2OS | Polyclonal | 74 |
| U2OS | Monoclonal 1 | 42 |
| A549 | Polyclonal | 90 |
| A549 | Monoclonal 1 | 98 |
| A549 | Monoclonal 2 | 93 |
| A549 | Monoclonal 3 | 94 |

Supplementary Table 1 - Measurement of CRISPR Efficiency in the clones depicted in Figure 3A. CRISPR Efficiency was measured by ability to remove expressed GFP from cells via FACS reporter assay - see Supplemental Methods section "Creating polyclonal and monoclonal Cas9 expressing cells lines and assessing Cas9 activity" for more information.

### Supplementary Methods

##### Design of the JUMP-MOA plate and plate layout

The JUMP-MOA list of compounds consists of 90 compounds derived from Broad Institute's Drug Repurposing Hub dataset[1]. We selected these compounds so that, in general, there are two compounds that belong to the same mechanism of action (MOA) class. We chose 90 compounds so that we could fit four replicates of each compound on a 384-well plate (plus many negative control wells).

To select these compounds, we began by filtering out compounds on the Organisation for the Prohibition of Chemical Weapons (OPCW) list of chemical weapons precursors or the U.S. Drug Enforcement Agency (DEA) list of controlled substances, from the repurposing hub compounds. We also excluded egregiously polypharmacologic compounds that are listed in the Chemical Probes Portal as “historical compounds”[2]. Additionally, we removed compounds annotated as targeting genes that are not expressed in U2OS or A549, as we performed all our experiments in either of these two cell types. Lastly, we filtered out those compounds without a sister compound belonging to the same MOA class.

To include confident positive pairs in our experiments, we selected pairs of chemical probes from the Chemical Probes Portal that are annotated as confidently having a particular MOA. We selected the remaining compounds from the filtered repurposing hub dataset. The 90 compounds we chose ended up belonging to 47 MOA classes; 43 MOA classes contain at least two compounds, while the other four contain only one compound (three instances of the same compound chosen twice whose similarity was not caught by our filters and one instance of compound unavailability).

We also included DMSO (Dimethyl sulfoxide) as the negative control, which is the solvent for the compounds. Thus, all wells contain the same concentration of DMSO, but the negative control wells do not have any additional compound treatment.

We designed the plate layout such that there are four replicates of each compound on the plate. This minimizes the influence of well-position effects, which causes treatments in the same well-position to correlate strongly with one another even if the treatments are dissimilar or inactive. We also have 24 negative control wells scattered across the plate, which can aid in the mitigation of well-position effects during profile normalization (Figure 2 A).

Additional information about these compounds is available at https://github.com/jump-cellpainting/JUMP-MOA. Note that eight of the 90 compounds were de-identified and are publicly identified only as *Compound1* through *Compound8* https://github.com/jump-cellpainting/JUMP-MOA#de-identified-compounds ).

The JUMP-MOA list of compounds and plate layout can be downloaded from https://github.com/jump-cellpainting/JUMP-MOA

##### Creating polyclonal and monoclonal Cas9 expressing cell lines and assessing Cas9 activity (CPJUMPA)

**Reagents not listed in main protocol**:

- pXPR_111 lentivirus (aka pLEX_311Cas9-v2, GPP vector link, internal reference lot #M-AE36-20190924)
- pXPR_047 lentivirus (GPP vector link, internal reference lot #M-AG54-20191217)
- Flow buffer (PBS, 2% FBS, 5 mM EDTA)
- PBS (Sigma-Aldrich Inc, cat. no. D8537-6X500ML)
- FBS (Sigma-Aldrich Inc, cat. no.F2442-500ML)
- EDTA, 0.5 M, pH 8.0 (Invitrogen, cat. no. 15575-038)

**Equipment not listed in main protocol**:

- Sterile 12-well plate multiple well plates (Corning, cat. no. 3512)
- T-75 tissue culture flask (Greiner bio-one, cat. no. 658170)
- V-bottom 96w plate (Greiner bio-one, cat. no. 651261)
- Flow cytometer (BD Biosciences, model Accuri™ C6 Plus Flow Cytometer)

**Reagent setup:**

- **Blasticidin selection** Added to growth media at a final concentration of 8 ug/mL for A549 and 16 ug/mL for U2OS **Puromycin selection** Added to growth media at a final concentration of 1 ug/mL
- **Flow cytometry buffer** 50 mL PBS + 1 mL FBS + 500 µLEDTA

Prior to running experimental batches with CRISPR knockout perturbations, cells stably expressing Cas9 must be generated and assessed for their Cas9 cutting efficiency. Parental A549 and U2OS cells were ordered from ATCC and maintained in culture using standard best tissue culture practices. The cells were then transduced with a lentiviral vector containing both a Cas9 expression cassette under an EF1a promoter and a blasticidin resistance cassette under an SV40 promoter. After transduction with the Cas9 vector, the cells were grown in media containing blasticidin to select for the Cas9-positive population. However, expression of Cas9 does not necessarily ensure that an adequate level of Cas9 cutting efficiency, i.e. the activity of the enzyme, will occur for CRISPR knockout. Therefore, a reporter assay can be used to assess the level of Cas9 activity prior to using the cells for downstream experiments. The reporter assay lentiviral vector contains an eGFP expression cassette, an sgRNA targeting eGFP and a puromycin resistance cassette. Cells without active Cas9 will express the eGFP and fluoresce, while cells with active Cas9 will cleave the eGFP and not fluoresce. The results of this reporter assay can therefore be easily read out via flow cytometry.

**Full list of experimental conditions**

1. A549 and U2OS cells were grown to near confluence (~80%) in a T-175 tissue culture flask then harvested and counted using a cell counter.
2. A master mix of cell suspension, Cas9 lentivirus (vector pXPR_111) and media were combined to a final volume of 22 mL. Polybrene was then added to the master mix at a final concentration of 4 µg/mL.
3. 2 mL per 12-well of the master mix was seeded into 11 wells of the 12-well plate such that each well contained 1.5x10e6 cells and 750 µL of Cas9 lentivirus.
4. A secondary master mix containing 1.5x10e6 cells, a final concentration of 4 µg/mL polybrene and up to 2 mL of media was mixed and seeded into the remaining well on the 12-well plate as the no-virus control.
5. The plate was then centrifuged for 2 hours at 1,000 x g at 30˚C and after the spin 2 mL per 12-well of fresh growth media was added dropwise to dilute the polybrene and lentivirus containing media. The plate was then moved into an incubator (37°C, 5% CO2).
6. The parental A549 and U2OS cells were maintained in culture as a control sample for the Cas9 activity reporter assay beginning in step 12.

7. 24 hours post-transduction, the cells were harvested out of the 12-well plate, and the cells that received the Cas9 lentivirus were pooled together separately from the no-virus control cells.
8. The two cell populations were counted, and each population was then seeded in duplicate into a 6-well plate at 150,000 cells per 6-well. The remaining Cas9-transduced cells were seeded into a T-175 tissue culture flask.
9. 24 hours post seeding, growth media containing blasticidin was added to one well of each cell population in the 6-well plate and to T175 flask at a final concentration of 8 ug/mL for A549 and 16 ug/mL for U2OS.
10. The no-virus control well that received blasticidin was monitored over the next several days to ensure that the cell population was fully selected.
11. The Cas9-transduced cells cultured in the T-175 flask were monitored and grown in blasticidin containing growth media for a total of 2 weeks post-transduction prior to proceeding with the Cas9 activity reporter assay.
12. The parental and polyclonal Cas9 A549 and U2OS cells were grown to near confluence (~80%) in a T-175 tissue culture flask, harvested and counted using a cell counter.
13. For both A549 and U2OS, 1.5x10e6 parental cells per well were seeded in duplicate 12-wells and 1.5x10e6 polyclonal Cas9 cells were seeded into one 12-well.
14. 200 µL of the Cas9 activity reporter assay lentivirus (vector pXPR_047) was then added to one of the duplicate parental 12-wells and the polyclonal Cas9 12-well.
15. Media up to 2 mL was added to all of the three 12-wells in addition to polybrene at a final concentration of 4 µg/mL.
16. The plate was then centrifuged for 2 hours at 1,000 x g at 30˚C and after the spin 2 mL per 12-well of fresh growth media was added dropwise to dilute the polybrene and lentivirus containing media. The plate was then moved into an incubator (37°C, 5% CO2).
17. 24 hours post-transduction, each cell population was harvested separately out of the 12-well plate and seeded into a T-75 tissue culture flask. Media containing puromycin was added to the parental and polyclonal Cas9 cells that were transduced with the reporter assay lentiviral vector at a final concentration of 1 ug/mL. The parental cells that were not transduced with the reporter assay were seeded in standard growth media only.
18. After five days, the three cell populations were harvested out of the T-75 flasks and the cell suspension was centrifuged for 5 minutes at 400 x 6 to pellet the cells.
19. The growth media was aspirated and the cells were resuspended in 1 mL of flow cytometry buffer. 200 µL of cell suspension in buffer was then added to one well of a 96-well V-bottom plate and the three cell populations were assayed by flow cytometry.
20. The parental cells that were not transduced with the reporter assay vector nor selected with puromycin were used to draw the appropriate gates for EGFP-negative cells. These same gates were used to assess the percentage of EGFP positive and negative cells for both the parental and polyclonal Cas9 cell populations that were transduced with the reporter assay vector.

##### Testing cell seeding densities, polybrene and selection drug doses at multiple time points across cell lines (CPJUMPB)

**Equipment not listed in main protocol**:
- 96-well Clear V-Bottom 2mL Polypropylene Deep Well Plate (Corning, cat. no. 3960)
- 96-well Clear V-Bottom Polypropylene Not Treated Microplate, Sterile (Corning, cat. no. 3357)

Prior to running experimental batches with either ORF overexpression or CRISPR knockout perturbations, we ran several initial experiments to finalize the conditions for transducing with lentiviral constructs. These included testing two readout time points for each perturbation, various cell seeding densities, and various polybrene and selection drug concentrations for eight different cell types (parental A549 and U2OS, polyclonal Cas9 A549 and U2OS, and monoclonal Cas9 A549 and U2OS). Each condition was assessed visually for cell confluency and health, an ideal confluency being 80% of the well surface area. Each cell plate was also analyzed for cell viability using CellTiter-Glo and a standard luminescence plate reader. The highest polybrene concentration that did not affect cell viability, and the lowest blasticidin or puromycin concentration that fully selected the cells, were chosen.

**Plate layout design**

The experiment was set up such that each cell line was seeded into one 384-well plate per time point and per perturbation type with varying cell seeding densities, polybrene and selection drug concentrations. For the plates mirroring the conditions to be used with ORF overexpression, parental A549 and U2OS were seeded at densities optimized for a readout time points of 48 and 96 h. For the plates mirroring the conditions to be used with CRISPR knockdown, polyclonal and monoclonal Cas9 expressing A549 and U2OS were seeded at densities for readout time points of 96 and 144 h. Four different cell densities were chosen for each cell line and seeded as six replicate columns per density. Per cell seeding density, three concentrations of polybrene and four concentrations of either blasticidin or puromycin were added. Each seeding density, polybrene and blasticidin or puromycin condition therefore had n=8 replicate 384-wells.

**Full list of experimental conditions**

1. A549 and U2OS cells were grown to near confluence (~80%) in a T-175 tissue culture flask then harvested and counted using a cell counter.
2. Each A549 and U2OS parental, polyclonal Cas9 and monoclonal Cas9 cell line was diluted to four varying concentrations based on prior calculations of the cell's doubling time and the chosen readout time point (A549 for 48h: 1425 - 2300 cells per 384-well; A549 for 96h: 250 - 725 cells per 384-well; A549 for 144h: 75 - 250 cells per 384-well; U2OS for 48h: 2250 - 3000 cells per 384-well; U2OS for 96h: 525 - 1400 cells per 384-well; U2OS for 144h: 175 - 775 cells per 384-well). The target cell confluency was 80% of the well surface area at the readout time point.
3. Each 384-well was then seeded at 40 µL in white, tissue culture-treated plates with clear flat bottoms using an automated liquid-handling system.
4. Cell plates were kept at RT for about 1h before proceeding.
5. During this 1h incubation period, separate stocks of growth media containing 0, 4, or 8 µg/mL polybrene were made. Each polybrene stock was then added to every third column of a 96-well plate.
6. Using an automated liquid-handling system, 10 µL of polybrene media was aspirated from the 96-well plate and added per 384-well such that the contents of each 96-well were dispensed into one quadrant of the 384-well plate.
7. The plates were then gently tapped to ensure an even distribution of cells across each well, and centrifuged for 30 minutes at 1,178 g and 37°C to mimic the centrifugation required during lentiviral transduction.
8. After centrifugation, the cell plates were placed in an incubator (37°C, 5% CO2) with alternating PBS plates to reduce edge effects.
9. 24 hours post-seeding, separate stocks of growth media containing blasticidin were made, either 0, 48, 56, or 64 µg/mL for the 48h readout time point or 0, 16, 32, or 48 µg/mL for the 96h readout time point. Separate stocks of growth media containing 0, 0.75, 1 or 1.5 ug/mL puromycin were also made. The varying concentrations of blasticidin and puromycin were then each added to two full rows of a 96-well plate.
10. 50 µL of media per 384-well was removed from the cell plates and replaced with 50 µL of media from the blasticidin or puromycin 96-well plates such that the contents of each 96-well were dispensed into one quadrant of the 384-well plate. Blasticidin was added only to the cell plates seeded with parental A549 and U2OS as they mirrored the conditions to be used with ORF overexpression, while puromycin was added only to the cell plates seeded with either polyclonal or monoclonal Cas9 A549 or U2OS as they mirrored the conditions to be used with CRISPR knockout. The cell plates were then returned to the incubator.
11. For the 144h readout time point cell plates only, a secondary media change was performed at 96h post-seeding by removing 50 µL of media and then replacing with 50 µL puromycin media as described in steps 9-10.
12. At each readout time point, plates were allowed to come to room temperature for 30 minutes before 10 µL per 384-well room temperature CellTiter-Glo was added. The plates were then covered with aluminum foil and put on a shaker at low speed for 15 minutes.
13. The plates were then imaged using a standard plate reader.

#### Testing cell seeding densities and lentivirus transduction volumes at multiple time points across cell lines (CPJUMPC)

**Reagents not listed in main protocol**:
- Control ORF 96w plate lentivirus (vector pLX_304, GPP internal plate identifier #DOG30)
- Control CRISPR 96w plate lentivirus (vector pXPR_003, GPP internal plate identifier #DXH92 for A549, #DXH94 for U2OS)

Prior to running experimental batches with either ORF overexpression or CRISPR knockout perturbations, we ran several initial experiments to finalize the conditions for transducing with lentiviral constructs. These included testing two readout time points for each perturbation, various cell seeding densities, and various volumes of the ORF overexpression and CRISPR knockout lentiviral constructs for four different cell types (parental A549 and U2OS and polyclonal Cas9 A549 and U2OS). Each set of conditions was plated in duplicate such that lentiviral transduction efficiencies could be calculated based on the number of cells surviving blasticidin or puromycin selection compared to the number of cells that did not receive selection drug treatment. Each condition was assessed visually for cell confluency and health, an ideal confluency being 80% of the well surface area.

Lentiviral transduction efficiencies and lentiviral toxicity was analyzed using CellTiter-Glo and a standard luminescence plate reader. The lentiviral volume that achieved a minimum transduction efficiency of 75% with minimal toxicity was chosen.

**Plate layout design**

The experiment was set up such that each cell line was seeded into one 384-well plate per time point and per perturbation type with varying cell seeding densities and lentiviral volumes. For the ORF overexpression plates, parental A549 and U2OS were seeded at densities optimized for a readout time points of 48 and 96 h. For the CRISPR knockdown plates, polyclonal Cas9 expressing A549 and U2OS were seeded at densities for readout time points of 96 and 144 h. Four different cell densities were chosen for each cell line and seeded as six replicate columns per density. These cell densities encompassed an ideal narrow range and were chosen based off of the results of the previous assay. Per cell seeding density, four concentrations of either ORF overexpression or CRISPR knockout lentivirus was added. Each seeding density and lentiviral volume condition therefore had n=48 replicate 384-wells.

**Full list of experimental conditions**

1. A549 and U2OS cells were grown to near confluence (~80%) in a T-175 tissue culture flask then harvested and counted using a cell counter.
2. Each A549 and U2OS parental and polyclonal Cas9 cell line was diluted to four varying concentrations (A549 for 48h: 1425 - 1500 cells per 384-well; A549 for 96h: 350 - 425 cells per 384-well; A549 for 144h: 100 - 110 cells per 384-well; U2OS for 48h: 1800 - 1875 cells per 384-well; U2OS for 96h: 400 - 475 cells per 384-well; U2OS for 144h: 135 - 150 cells per 384-well). The target cell confluency was 80% of the well surface area at the readout time point.
3. Each 384-well was then seeded in duplicate at 40 µL in white, tissue culture-treated plates with clear flat bottoms using an automated liquid-handling system. Cell plates were kept at RT for about 1h before proceeding.
4. During this incubation period, 96-well plates containing ORF overexpression or CRISPR knockout lentivirus were thawed on wet ice, then quickly pulse spun to prevent any virus from clinging to the plate seals. The plates were placed back on ice until needed.
5. Then, using an automated liquid-handling system, 10 µL of polybrene media was added per 384-well plate (final concentration 4 ug/mL).
6. Immediately after polybrene addition, lentivirus was aspirated from the 96-well plate and added per 384-well such that the contents of each 96-well were dispensed into one quadrant of the 384-well plate at volumes of 0, 0.5, 1 and 2 µL.
7. The plates were then gently tapped to ensure an even distribution of cells across each well, and centrifuged for 30 minutes at 1,178 g and 37°C.
8. After centrifugation, the cell plates were placed in an incubator (37°C, 5% $CO_2$) with alternating PBS plates to avoid edge effects.
9. 24 hours post-seeding, 50 µL of media per 384-well was removed from the cell plates. For one of the duplicate cell plates, the media was replaced with 50 µL per 384-well of fresh growth media. For the other duplicate cell plate, the media was replaced with 50 µL per 384-well of media containing either 16 µg/mL blasticidin or 0.75 µg/mL puromycin. Blasticidin was added only to the cell plates seeded with parental A549 and U2OS, while puromycin was added only to the cell plates seeded with either polyclonal Cas9 A549 or U2OS. The cell plates were then returned to the incubator.
10. For the 144h readout time point cell plates only, a secondary media change was performed at 96h post-seeding by removing 50 µL of media and then replacing with either 50 µL fresh growth media or 50 µL puromycin media as described in step 9.
11. At each readout time point, 10 µL per 384-well room temperature CellTiter-Glo was added. The plates were then covered with aluminum foil and put on a shaker at low speed for 15 minutes.
12. The plates were then imaged using a standard luminescence plate reader.

### Supplementary References


1. Corsello, S. M. *et al.* The Drug Repurposing Hub: a next-generation drug library and information resource. *Nat. Med.* **23**, 405–408 (2017).

2. Arrowsmith, C. H. *et al.* The promise and peril of chemical probes. *Nat. Chem. Biol.* **11**, 536–541

(2015).
